# Dietary depletion of glutamine is atheroprotective

**DOI:** 10.64898/2026.03.06.710174

**Authors:** Anita Salamon, Pernilla Katra, Victoria M. Milosek, Rupa Tripathi, Ria Raval, Laura Shankman, Alexandra Krinsky, Noelle Koo, Sohel Shamsuzzaman, Daniel Engelbertsen, Eva Bengtsson, Xiaoke Yin, Honglin Chen, Stefan Bekiranov, Manuel Mayr, Harry Björkbacka, Vlad Serbulea, Gary K. Owens

## Abstract

Heart attacks and strokes are late-stage complications of rupture of unstable atherosclerotic plaques. Stable plaques contain stabilizing matrix-producing fibrotic cells, largely smooth muscle cell (SMC)-derived. The molecular drivers of SMC phenotypic transitions to beneficial fibrotic or destabilizing inflammatory and calcifying phenotypes are unclear. Since atherosclerosis develops over decades, there is extensive interest in identifying dietary alterations that enhance plaque stability. We demonstrate that SMC acquire a fibrotic phenotype dependent on glutamine-derived metabolites supporting both catabolism and collagen synthesis. Moreover, dietary glutamine restriction decreases mortality of mice susceptible to atherosclerotic plaque rupture. Lesions from glutamine-restricted mice are smaller and have increased SMC investment. This study identifies dietary glutamine as a driver of cardiovascular mortality, suggesting a new strategy for reducing late-stage complications of atherosclerosis.

## Main Text

Clinical manifestations of vulnerable atherosclerotic plaques, such as myocardial infarction (MI) and stroke, are the leading cause of death in the USA and worldwide ^1^. MI and stroke arise from rupture of atherosclerotic plaques ^2^. Plaque rupture physically disrupts the fibrous cap (FC) ^3^, a protective extracellular matrix (ECM)-rich layer overlying atherosclerotic lesions, exposing pro-thrombotic lesion matter to circulating platelets which initiate intramural clot formation. Current treatments of advanced atherosclerosis have focused on aggressive lipid lowering primarily with statins ^4^ and PCSK9 inhibitors alongside lifestyle and dietary modifications. The latter has the potential advantage of safety, early intervention, and long-term efficacy in inhibiting development of atherosclerosis rather than the more difficult process of regressing lesions ^5^. Aside from the notable benefits of reducing dietary lipids and sugar, we have a poor understanding of dietary mechanisms affecting lesion pathogenesis and have not identified targets for inhibiting lesion development and progression despite over six decades of research. Central to the stability of advanced atherosclerotic plaques is the presence of a thick FC containing predominately collagens and proteoglycans produced by smooth muscle cells (SMC) that have transitioned to a fibrotic phenotype (SMC-F) ^6,7^. Studies from our lab and others using murine models of atherosclerosis (*Apoe*^-/-^) combined with SMC-specific lineage tracing (*Myh11*-CreER^T2^ <STOP>^FL/FL^-*eYFP*^+/+^) and knockout of genes such as *Klf4*, *Oct4*, *Pdgfrb*, and *Col15a1* have shown that SMC phenotypic transitions play a critical role in the development of atherosclerosis ^8–11^.

It was initially assumed that SMC phenotypic transitions to a proliferative state would be detrimental for lesion stability ^12^, establishing a two-state system of beneficial versus detrimental SMC. However, single cell RNA sequencing (scRNA-seq) analyses of diseased arteries by our lab and others have shown that SMC transition to multiple phenotypes ^8,13–16^, whose effects on plaque stability are presumed based on their inherent transcriptional profiles. Some studies identified foam cells and osteoblasts derived from SMC, thought to be inflammatory and calcifying respectively, but all studies show evidence for SMC assuming a dedifferentiated stem-cell-like state, referred to as synthetic SMC, and transitioning to a fibrotic ECM-remodeling state ^8,13–16^, referred to as SMC-F from here on out. As such, a major focus is to identify factors and mechanisms that promote beneficial (i.e. plaque-stabilizing) changes in SMC phenotype, such as that of SMC-F. Our lab and others have shown that SMC to SMC-F transitions require PDGF and TGFB signaling ^8,17–20^.

Recent studies have revealed that stem cells tend to reside in hypoxic niches and, like some tumor cells, heavily depend upon lactate-producing *aerobic glycolysis* for proliferation. We have previously demonstrated that SMC-F transitions are dependent upon aerobic glycolysis ^8^. Further characterization of stem-cell metabolism revealed a dependency on glutamine (Gln), the most abundant dietary amino acid. Gln has several identified functions within a cell, including serving as a catabolic substrate via alpha-ketoglutarate and serving as an anabolic source of proline for protein synthesis. Metabolomic analysis of human carotid endarterectomies revealed that, in addition to glucose and lactate, Gln was significantly changed between stable and unstable plaques ^21^. Few studies have investigated the role of Gln in atherosclerosis. One study used both adenovirus-based knockout and genetic deletion of glutaminase 2 (*Gls2*) in atherosclerotic *Apoe*^-/-^ and *Ldlr*^-/-^ mice ^22^. Perturbation of *Gls2* led to augmented levels of Gln in plasma, which in turn was associated with enhanced plaque burden and lesion size. Conversely, overexpression of *Gls2* had the opposite effect – lowering plasma Gln and decreasing lesion size. Although this study addressed whether Gln affects plaque size, it did not address whether it plays a role in plaque stability and risk of cardiovascular events. Furthermore, it is not known if SMC phenotype is influenced by Gln availability nor whether dietary Gln manipulation can influence plaque stability.

## Results

### Gln is required for the transition of cultured SMC to a fibrotic state

To address these questions, we first tested whether Gln availability plays a role in an established model of SMC-phenotypic switching *in vitro* ^8^. Here we cultured SMC lines from the thoracic aorta of SMC lineage-tracing mice, which upon growing to confluence are switched to serum-free media (SFM). These SMC adopt a contractile phenotype (SMC-C) characterized by expression of contractile genes such as *Myh11*, *Acta2*, and *Cnn1* ^23,24^. SMC-C treated with recombinant murine platelet-derived growth factor subunit B homodimer (PDGF) and transforming growth factor beta 1 (TGFB) transitioned to a fibrotic state (SMC-F; Fig. 1A), defined namely by: upregulation of fibrotic genes (fig. S1, A to D), major changes in morphology, augmentation of migration and proliferation (fig. S1, E to G), and changes in cellular metabolism (fig. S1, H and I). Expanding these studies, bulk RNA sequencing (RNA-seq) and gene-set enrichment analysis of SMC-F highlights a matrix-producing phenotype with positive enrichment for *Extracellular Matrix Organization (ECM)* related pathways (fig. S1A, and data S1). Leading-edge analysis revealed 16 markers that drive the enrichment of multiple ECM gene sets including *Col15a1*, *Fn1*, *Itgb3*, and *Spp1*, highlighting their central role in the matrix-remodeling program adopted by SMC-F (fig. S1, A to D, and data S1).

**Fig. 1.**
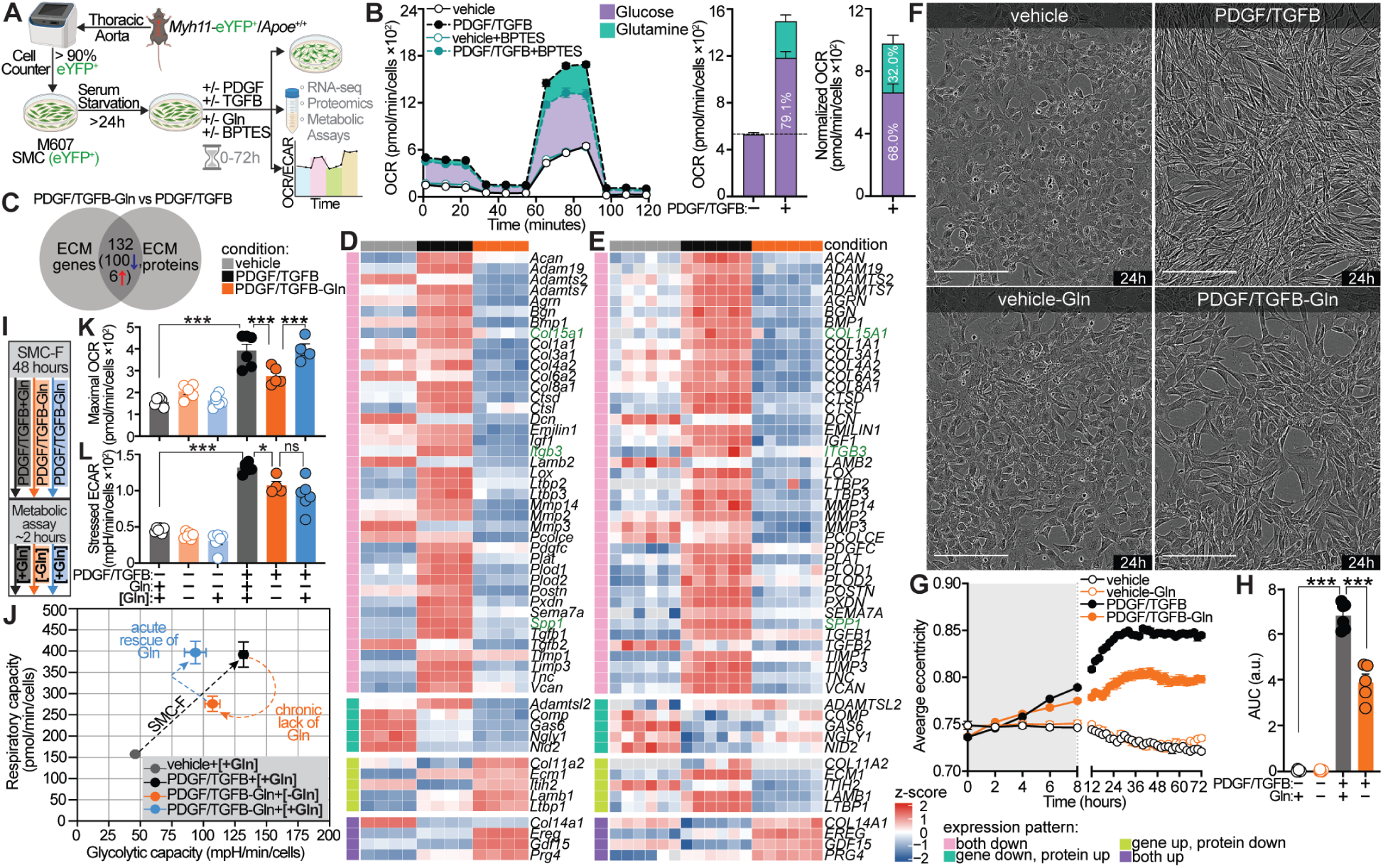
Glutamine is required for SMC-F transition in vitro. (**A**) Experimental design. Aortic smooth muscle cells (SMC; line M607) were plated and serum-starved prior to treatment with or without 30 ng/mL PDGF-BB (PDGF), 10 ng/mL TGFB, Gln (2 mM), and glutaminase inhibitor BPTES (5 μM). (**B**) Mitochondrial stress test measuring oxygen consumption rate of SMC with or without BPTES. The substrate contribution to the maximum respiratory capacity (over vehicle control) of the PDGF/TGFB conditions is shown (middle and right; SMC-F OCR). (**C**) Venn diagram showing co-occurrence of significantly expressed ECM proteins and genes in PDGF/TGFB-Gln compared to PDGF/TGFB treated cells after 24 hours (*P* < 0.05). Of a total of 132 ECM genes in common between the groups, 75% were downregulated in the absence of Gln. A complete list of common ECMs is presented in data S2. Heatmap of ECM (**D**) genes or (**E**) proteins (adjusted *P-*value < 0.05), grouped by expression pattern between conditions. Red indicates increased expression, blue indicates decreased expression, and white indicates no change. In green, ECM-markers of SMC-F were highlighted. (**F**) Representative phase contrast images after 24 hours of treatment. Scale bar, 200 μm. (**G**) SMC morphology was measured every 2 hours for 3 days. Each data point represents the average of 4-8 technical replicates. Grey shading highlights a Gln-independent phase of SMC eccentricity. (**H**) Quantification of cell eccentricity after 72 hours (a.u.: arbitrary unit). (**I**) Schematic of M607 treated with vehicle, PDGF/TGFB, and/or Gln for 48 hours. Respiratory and glycolytic capacities were then assessed using media with or without Gln (highlighted in bold [+/- Gln]). (**J**) Energy capacity map representing the bioenergetic potential of SMC using glycolytic capacity (x-axis; stressed ECAR) and respiratory capacity (y-axis; maximal OCR). (**K**) Gln supplementation after extended Gln-withdrawal, rescued respiration, but not (**L**) glycolysis in SMC-F. Graphs were analyzed using one-way ANOVA with Tukey’s correction for post-hoc analysis with N ≥ 3, error bars represent mean ± SEM. \*\*\**P* < 0.001; \*\**P* = 0.001 to 0.01; \**P* = 0.01 to 0.05; ns, not significant.

We have previously shown that SMC-F depend upon rerouting glucose metabolism from the TCA cycle and towards lactate production to support ECM synthesis ^8^. Despite this metabolic rewiring, SMC treated with PDGF and TGFB had an increased oxygen consumption rate (OCR), a measurement for oxidative phosphorylation and TCA cycle utilization (fig. S1, H and I). We hypothesized that SMC-F must adopt a catabolic substrate in addition to glucose to sustain the TCA cycle. To determine if Gln serves as a catabolic substrate in cultured SMC, we quantified mitochondrial respiratory capacity, the maximum potential of cells to use oxidative phosphorylation and the TCA cycle, using extracellular flux analysis (Fig. 1, A and B). We found that after 24 hours, 32% of OCR in SMC-F cells are Gln-dependent (Fig. 1B). This data suggests that unlike contractile SMC, SMC-F catabolize Gln.

To test whether Gln is necessary for the SMC-F transition, we cultured SMC with or without Gln in the presence of PDGF and TGFB (Fig. 1A). SMC treated in the absence of Gln had significantly reduced eccentricity, a less-spindly morphology (Fig. 1, F to H), and inhibited ECM gene and protein expression (Fig. 1, C to E, and data S2), most likely due to metabolic insufficiency for biosynthesis. We developed an analytic method using the Incucyte Cell Analysis software where individual cells can be classified as having a morphology consistent with either SMC-C or SMC-F (fig. S2). SMC without Gln during PDGF and TGFB treatment failed to increase the glycolytic and respiratory capacities characteristic of SMC-F (Fig. 1, I to K). Acute reintroduction of Gln to SMC-F that lacked Gln for 48 hours rescued respiratory but not glycolytic capacity (Fig. 1, I to K). This demonstrates that Gln presence throughout the phenotypic transition is at least partially required for the increase in glycolytic capacity.

However, despite the contribution of Gln to respiration in SMC-F, it is not required for the augmentation of respiratory capacity during the transition – only at the end. Similarly, SMC eccentricity changes indicate that the first 8 hours after PDGF and TGFB treatment do not require Gln (Fig. 1G). These data indicate that SMC depend upon Gln to undergo phenotypic transitions, but it is unclear if Gln plays a role in the initial dedifferentiation or only later during ECM synthesis.

### Gln is temporarily dispensable for SMC dedifferentiation, migration, and proliferation

To test whether Gln is required for SMC dedifferentiation into a proliferative and migratory state prior to fibrosis, we performed a scratch wound assay over a 96-hour treatment period, which depends on both proliferation and migration for wound closure. Additionally, we conducted bulk RNA-seq and extracellular flux analysis to profile gene expression and energy metabolism of these SMC (Fig. 2A). Principal component analysis (PCA) revealed substantial transcriptional changes in response to PDGF and TGFB as early as 4 hours (Fig. 2B). Unexpectedly, Gln was dispensable for the gene expression changes seen within the first 8 hours of treatment (Fig. 2, B to K; figs. S3 and S4; data S3). Within this 8 hour time period, wound closure (Fig. 2, C to E; fig. S3A), expression of migration-associated *Signaling by Rho GTPases* and *Signaling by GPCR* pathways and genes (Fig. 2F; fig. S4, A and C; data S3), and expression of pathways and genes linked to *ECM Organization*, *Signal Transduction*, and *Metabolism* (Fig. 2G) were unaffected by the absence of Gln. Beyond 8 hours however, Gln absence significantly impaired wound closure (Fig. 2, C to E). By 24 hours, expression of *ECM Organization*- and *Signal Transduction*-related genes and pathways, including *Signaling by Rho GTPases*, was markedly reduced in Gln-deprived SMC (Fig. 2, F and K; fig. S4, A and C). Given the established roles of Rho GTPases in cell cycle progression, proliferation, actin polymerization, and motility, their transcriptional suppression provides a mechanistic explanation for the observed reduction in migration.

**Fig. 2.**
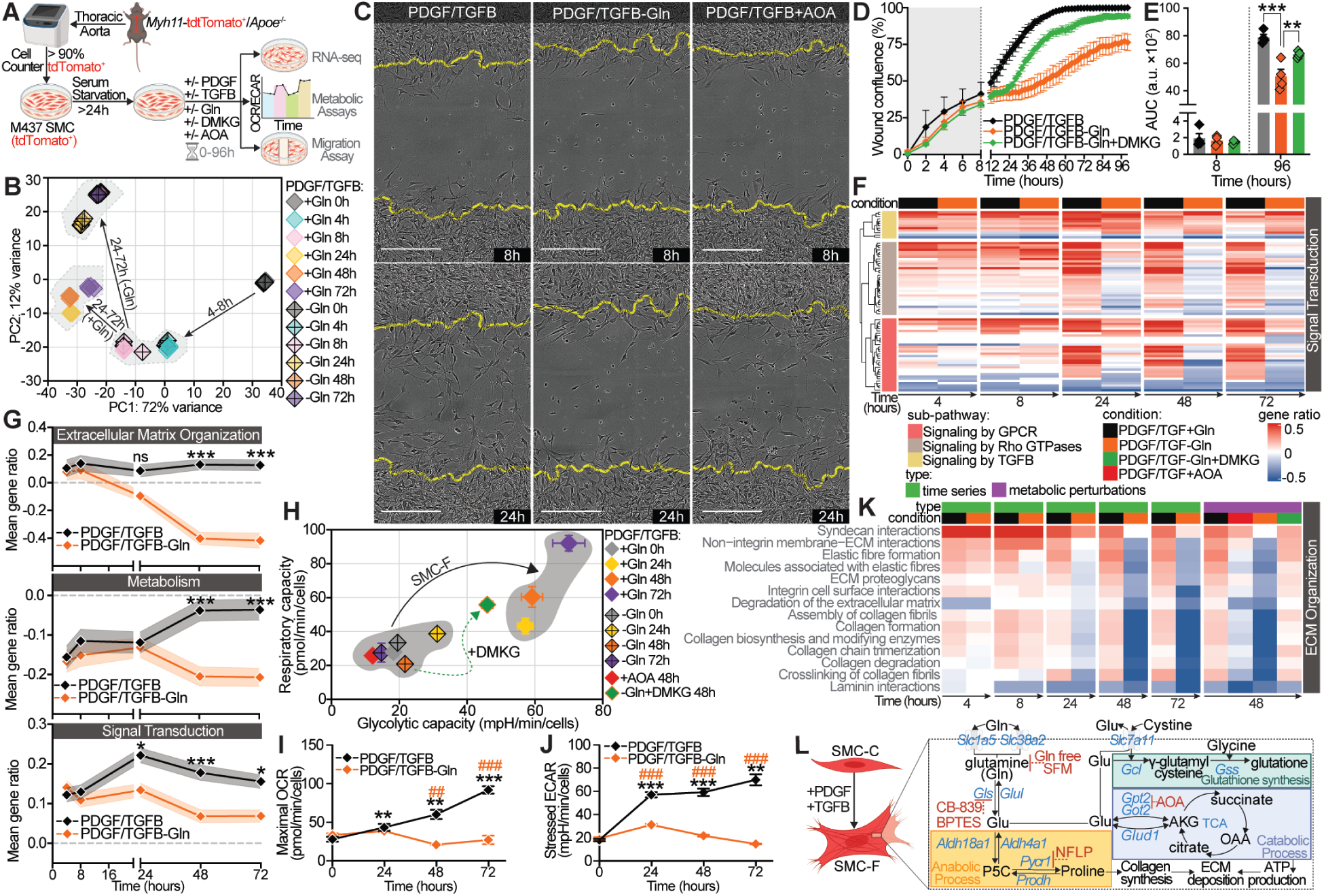
PDGF/TGFB-induced SMC phenotypic transitions become dependent on Gln after 8 hours. (**A**) Experimental design: SMC (M437, cultured from *Myh11*-Cre/tdTomato^+/+^/Apoe^-/-^male mice) treated with PDGF/TGFB ± glutamine (Gln), DMKG, or aminooxyacetate (AOA). (**B**) Principal component analysis of bulk RNA-seq transcriptional changes. (**C**) Representative migration assay images of SMC at 8- and 24-hours post-treatment. Yellow line indicates the initial scratch borders. Scale bar: 200 µm. (**D**) Wound confluence over time, and (**E**) area under the curve after 8 and 96 hours. Grey shading highlights a Gln-independent wound closure phase. (**F**) *Signal Transduction* super-pathway heatmap hierarchically clustered within sub-pathways over time, showing Gln- and time-dependent effect in *Rho GTPases* and *GPCR signaling.* (**G**) Effect of glutamine depletion on super-pathway expression pattern over time. (**H** to **J**) Bioenergetic capacity of SMC treated with PDGF/TGFB ± Gln, AOA, or DMKG over time. Statistical indicators: *PDGF/TGFB vs 0-hour; #PDGF/TGFB-Gln vs PDGF/TGFB+Gln. (**K**) Super-pathway analysis of *ECM Organization* (left) over time, and (right) with metabolic perturbations at 48 hours. (**L**) Schematic demonstrating contribution of Gln to metabolic pathways in SMC-F. Gln transporters*: Slc1a5*, *Slc38a2*; *Glud*, glutamate dehydrogenase; *Gpt*, glutamic-pyruvate transaminase; *Got*, glutamic-oxaloacetic transaminase. *Gcl*, glutamate-cysteine ligase; *Gss*, glutathione synthetase. In (F), and (K) rows represent individual pathways. Positive gene ratios (red) indicate net upregulation and blue indicates net downregulation relative to 0-hour. For (F), (G) and (K) see materials and methods for super-pathway analysis and data S3-S4. Statistics: two-way ANOVA with Sidak’s or (E) one-way ANOVA with Tukey’s correction. Error bars represent mean ± SEM, *N* ≥ 4. \*\*\**P* < 0.001; \*\**P* = 0.001 to 0.01; \**P* = 0.01 to 0.05; ns, not significant.

Gene expression associated with *Metabolism* is consistent through 24 hours of treatment, but fails to increase by 48 hours, as can be seen in the Gln-containing controls (Fig. 2G and data S3). The *Metabolism* gene expression is consistent with the functional metabolic readout, measured by extracellular flux analysis, showing that SMC treated with PDGF and TGFB maintain respiratory capacity for the first 24 hours irrespective of Gln presence (Fig. 2H). After 24 hours of treatment, only SMC treated in the presence of Gln increase respiratory capacity (Fig. 2, H and I). However, glycolytic capacity of SMC lacking Gln failed to increase after treatment with PDGF and TGFB (Fig. 2, H and J).

These data suggest that Gln is necessary for SMC to clear a checkpoint, allowing for dedifferentiation, migration, and proliferation. *In vitro*, this checkpoint occurs sometime between 8 and 24 hours after PDGF and TGFB treatment. The question remains as to whether it is Gln itself or a downstream metabolite that is used by SMC to support dedifferentiation, migration, and proliferation.

### Gln-derived metabolites provide both energy and substrates for ECM synthesis in SMC-F

Gln participates in several metabolic pathways (Fig. 2L). Gln can be deaminated, resulting in ammonia and glutamate (Glu). The ammonia generated during deamination of Gln can participate in glycosaminoglycan synthesis, needed for proteoglycan production – key components of ECM. Glu can be further deaminated to alpha-ketoglutarate (AKG) to enter the TCA cycle and contribute to ATP production ^25,26^. Glu can be converted to proline (Pro) to support protein synthesis through pyrroline-5-carboxylate synthase (*Aldh18a1*) and pyrroline-5-carboxylate reductase (*Pycr1*) ^27,28^. Additionally, Glu is an obligatory substrate for the synthesis of glutathione, serving to control redox balance within cells.

Initially, we found that SMC treated with PDGF and TGFB significantly induces the expression of genes involved in the Gln-to-Pro conversion pathway, including *Aldh18a1, Pycr1*, *Pycr2*, and *Pycr3* (fig. S5B), suggesting that SMC metabolize Gln to Pro to, in turn, synthesize Pro-rich collagens. To test if Gln-derived Pro supports collagen synthesis in SMC-F, we inhibited *Pycr1* with N-Formyl-L-proline (NFLP) ^29^ (fig. S5A). Co-treatment of SMC with PDGF and TGFB and NFLP resulted in no significant differences in ECM-gene expression, migration and proliferation, or energy metabolism (fig. S5, C to F). Interestingly, transcription and secretion of Pro-rich collagens, including Collagens VIII (*Col8a1*) and IV (*Col4a1*) whose amino acid composition is over 20% Pro ^30^, were significantly reduced in SMC treated with NFLP (fig. S5, G and H). These data suggest that Gln-derived Pro is essential for SMC-F collagen generation, but not other measured properties and functions of SMC-F.

Next, we asked whether Gln supports dedifferentiation and migration by serving as a substrate for ATP synthesis through its downstream metabolite AKG. To answer this question, we co-treated SMC with PDGF and TGFB and AOA, which is an inhibitor of pyridoxal phosphate dependent enzymes, such as GPT2 and GOT2 – an enzyme that converts glutamate to AKG (Fig. 2L) ^31,32^. Much like SMC in the absence of Gln, AOA-treated SMC had a similar defect in migration and proliferation (Fig. 2C, and fig. S3). Next, we asked whether the Gln-AKG axis is necessary for SMC transition to a fibrotic state. Bulk RNA-seq shows that SMC treated with PDGF and TGFB and AOA for 48 hours had inhibited expression of *ECM Organization* (Fig. 2K), *Metabolism* (fig. S4B), and *Signal Transduction* super-pathways, compared to control SMC (fig. S4A). SMC treated with PDGF and TGFB and AOA failed to augment either respiratory or glycolytic capacity by 48 hours, suggesting that the conversion of Gln to AKG is necessary for SMC phenotypic switching (Fig. 2H).

To test this hypothesis, we treated SMC with dimethyl alpha-ketoglutarate (DMKG), a cell-permeable analog of AKG, in the presence of AOA or in the absence of Gln – both conditions which result in defective migration and proliferation in a scratch wound assay (Fig. 2D and fig. S3). In the presence of the aminotransferase inhibitor AOA, DMKG completely rescued the PDGF and TGFB induced migration and proliferation of SMC (fig. S3). However, in the absence of Gln but with AKG, the migratory defect is partially, but not fully, rescued (Fig. 2, D to E; fig. S3). Similarly, we find that some, but not all, *Signal Transduction* pathways are rescued by AKG in the absence of Gln (fig. S4A and data S4). After 48 hours, all but one *Extracellular Matrix Organization* pathway was completely ablated in the absence of Gln and rescued by AKG (Fig. 2K). The *Signal Transduction* pathways, namely *Signaling by GPCR* and *Signaling by Rho GTPases* were inhibited in the absence of Gln (fig. S4C and data S4). These pathways were only partially rescued in the presence of AKG (fig. S3E). Expression of *Metabolism* pathways were substantially inhibited in the absence of Gln (fig. S4, B and D), but, surprisingly, not rescued by the presence of AKG (fig. S4, B and F and data S4). In the absence of Gln, AKG nearly completely restored energy production capacities, as measured by extracellular flux analysis (Fig. 2H). These data demonstrate that Gln feeds into the TCA cycle, at least in part through its metabolite AKG, and is required for SMC to phenotypically transition to a fibrotic state.

Taken together, these results suggest that Gln-derived AKG supports cellular programming responsible for ECM production but not metabolic reprogramming. Furthermore, these data show that the conversion of Gln to AKG is necessary but not sufficient for the SMC-F transition. Furthermore, it appears that Gln has an earlier role in SMC phenotypic switching.

### Gln is required for SMC phenotypic switching in vivo

Given the importance of SMC phenotypic switching during atherosclerosis, we hypothesized that Gln levels can be manipulated to modify atherosclerotic plaque stability. We and others have previously associated the cellular composition of plaques with their likely stability. To better understand how dietary Gln affects lesion cellular composition, we performed a series of transcriptomic analyses. First, we used bulk RNA-sequencing on brachiocephalic arteries of SMC-lineage tracing atherosclerotic mice (*Myh11-*Cre*/*tdTomato*^+/+^/Apoe^-/-^*) fed a Gln-free WD or Ctrl WD for 18 weeks (Fig. 3A). Pathway analyses revealed that contractile programming was upregulated in Gln-free mice (Fig. 3B). However, this analysis does not allow us to understand how relative proportions of SMC phenotypes are changed within atherosclerotic vessels. To address this question, we used scRNA-seq to characterize the SMC-derived cell phenotypes present within atherosclerotic brachiocephalic arteries (Fig. 3, A and C). Our analysis identified 5 unique SMC-derived (tdTomato^+^) clusters (Fig. 3, D to F) which we annotated as contractile (SMC-C1, SMC-C2), transitioning (SMC-T), and fibrotic (SMC-F1, SMC-F2) (Fig. 3F). The SMC in our chow-fed control mice, which did not develop noticeable atherosclerotic lesions, were almost exclusively SMC-C1 and SMC-C2 (Fig. 3I), both of which are enriched in contractile gene expression (Fig. 3J and fig. S6H; and data S5). Relative to SMC-C1, SMC-C2 appears to have a reduced metabolic gene expression (Fig. 3K and fig. S6, A to C), and an enhancement in *Signaling by Rho GTPase* genes (fig. S6, D to E; and Fig. 3K: *Signal Transduction*), suggesting enhanced motility and/or proliferative ability. After 18 weeks of Western diet feeding, we found significant proportions of SMC adopted a transitioning (SMC-T) or fibrotic (SMC-F) phenotype (Fig. 3, H and I).

**Fig. 3.**
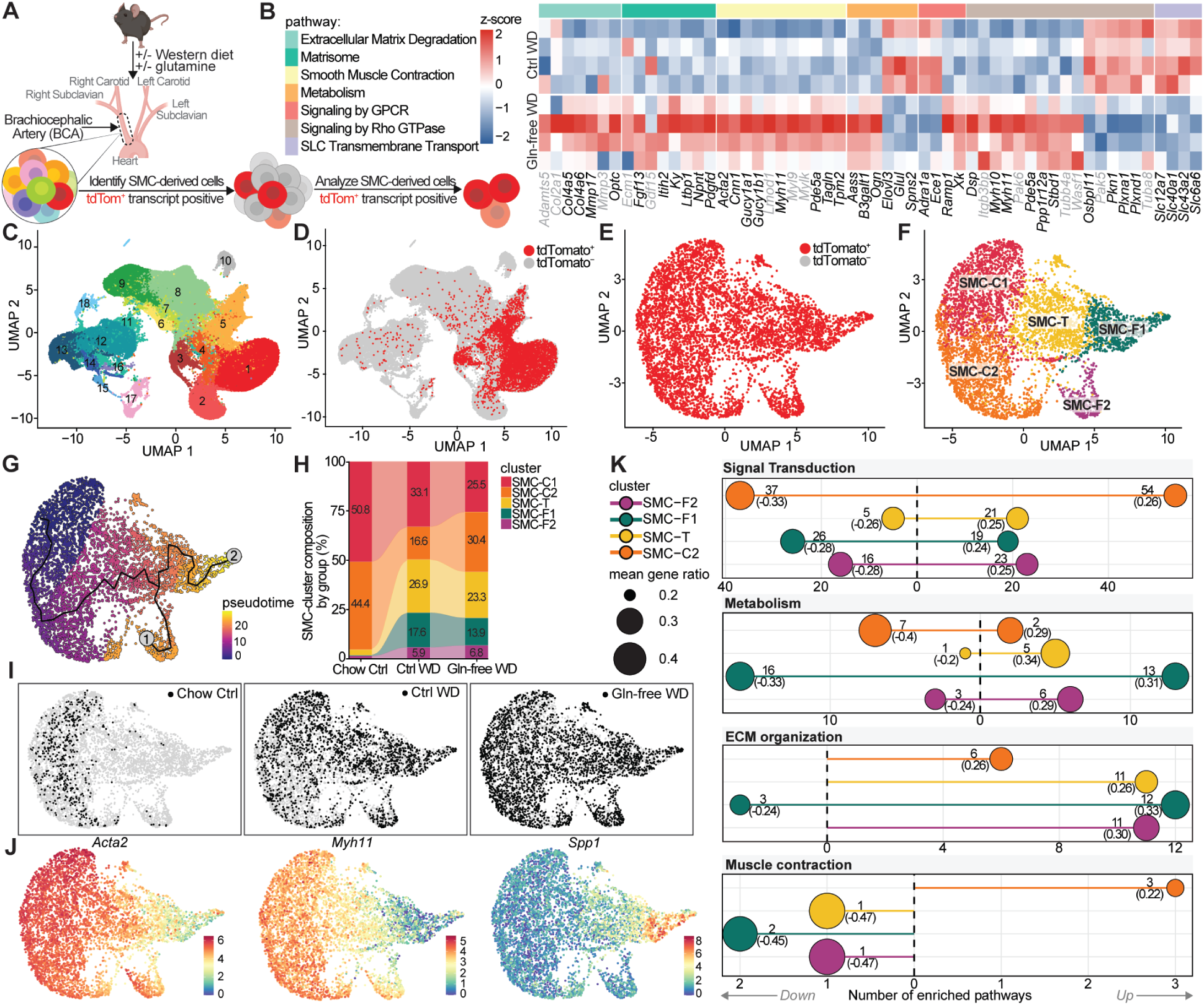
Glutamine is required for SMC phenotypic switching in vivo. (**A**) Schematic of scRNA-seq and bulk RNA-seq experimental design. (**B**) Heatmap of differentially expressed genes (adjusted *P-*value < 0.05) from bulk RNA-seq analysis of BCA grouped by Reactome pathways. Genes in grey denote genes of interest that were not statistically significantly changed. *N* = 4. (**C**) UMAP of 98,602 cells from all libraries. (**D**) tdTomato expression across all cells (9,642 tdTomato^+^ cells), and (**E**) tdTomato^+^ cells from male samples (5,636 tdTomato^+^ cells). (**F**) Subsetting and clustering of tdTomato^+^ cells from male mouse samples. (**G**) Pseudotime trajectory analysis of SMC phenotypic transitions. Terminal points of trajectory branches are indicated by numbered nodes. (**H**) Quantification of each SMC cluster from each sample group. (**I**) UMAP representing contribution of each sample group. (**J**) Feature plots of selected phenotypic markers. (**K**) Lollipop plots showing the number of enriched pathways within each Reactome super-pathway (x-axis) – the annotations indicate *#pathways (gene ratio)*. Pathways in super-pathway analysis were filtered for ≥ 10 pathway size, ≥ 2 DE genes, and |gene ratio| ≥ 0.2 in at least one direction (up- or downregulated). Mean gene ratio was calculated as the average of all up- and downregulated gene ratios. Data derived from differential expression analysis focusing on genes expressed by more than 50% of cells and with a log_2_(FC) ≥ 0.25 in 3 samples. FC: fold change. Some genes are annotated to multiple Reactome pathways.

To better understand how SMC phenotypic transitions progress in atherosclerotic lesions, we used Monocle 3 pseudotime analysis ^33,34^. This analysis allows us to postulate a differentiation path using the gene expression profiles of each cell, without explicit time-varied data. Based on this analysis, we postulated that classically-defined contractile SMC (SMC-C1) progress through two intermediates (SMC-C2 and SMC-T) prior to becoming one of two fibrotic phenotypes (SMC-F1 and SMC-F2) (Fig. 3G; and fig. S6). As described above, SMC-C2 cells have a transcriptomic profile consistent with a proliferative and migratory cell while maintaining contractile programming. Following SMC-C2, the SMC-T cells have lowered expression of contractile genes, instead expressing genes like *Ly6a* and *Lgals3*, consistent with a fully dedifferentiated stem cell-like phenotype (fig. S6F). From SMC-T, our analysis suggests that two distinct phenotypes can be achieved, annotated here as SMC-F1 and SMC-F2. Both SMC-F1 and SMC-F2 have enhanced ECM-gene expression (Fig. 3, J and K; fig. S6G), suggesting that these SMC transitioned to a fibrotic phenotype. Interestingly, we found that SMC-F2 has not only enhanced fibrotic expression but also maintains a substantial level of contractile gene expression, such as *Acta2*, *Myh11, and Cnn1* (Fig. 3J; and fig. S6H). These SMC-F2 cells may represent SMC that have transitioned through a fibrotic phenotype in the fibrous cap and then began reverting to a contractile state. Alternatively, these cells may represent a distinct myofibroblast-like phenotype, which share both fibrotic and contractile ability ^35,36^. Of note, the proportion of these transcriptional populations are different between Ctrl WD and Gln-free WD (Fig. 3H). The two sample groups had similar proportions of SMC-C1 (33.1% in Ctrl WD; 25.5% in Gln-free WD) and SMC-F2 (5.9% in Ctrl WD; 6.8% in Gln-free WD), a mild-to-modest decrease in SMC-T (26.9% in Ctrl WD; 23.3% in Gln-free WD) and SMC-F1 (17.6% in Ctrl WD; 13.9% in Gln-free WD) and a doubling of SMC-C2 (16.6% in Ctrl WD; 30.4% in Gln-free WD). These data suggest that, in the absence of Gln, the balance of SMC phenotypic states shifts to favor SMC-C2 – a migratory phenotype as indicated above. These data reveal a possible checkpoint in the process of phenotypic switching prior to transitioning to stem-like (SMC-T) and fibrotic states (SMC-F1).

In other words, our data suggest that Gln availability may not only affect the terminal fibrotic phenotype as we initially thought, but also plays a much earlier role in regulating SMC de-differentiation and migration into lesions. This is supported by data showing that loss of dietary Gln resulted in a greater proportion of SMC having a contractile phenotype (Fig. 3H).

Considering that, historically, human lesion stability was correlated with presence of ACTA2-positive cells ^37^ – likely contractile SMC – our findings predict that loss of dietary Gln would be plaque-stabilizing.

### Atherosclerotic mice fed a Gln-free diet experienced notable benefits in indices of plaque stability compared to control-diet fed mice

We hypothesized that Gln restriction, which increases the proportion of SMC-C within lesions (Fig. 3), would improve lesion stability. We then asked whether reduction of Gln intake could reduce atherosclerotic plaque formation. To answer this question, we fed *Myh11*-Cre/tdTomato^+/+^/*Apoe*^-/-^ mice ^38^ a tamoxifen-containing diet from 6 to 8 weeks of age to irreversibly lineage tag SMC. At 9 weeks of age, mice were fed WD with and without Gln (Ctrl WD/Gln-free WD, respectively) for 18 weeks to form advanced atherosclerotic lesions (Fig. 4A). We found that Gln-free WD fed mice had smaller brachiocephalic artery (BCA) lesions, greater lumen area, and greater medial thickness, as determined by MOVAT staining (Fig. 4C). This occurred despite the Gln-free WD causing no changes in plasma lipids or liver enzymes, albeit there was a noted decrease in heart and lung weight in these Gln-free WD-fed mice (Fig. S7).

**Fig. 4.**
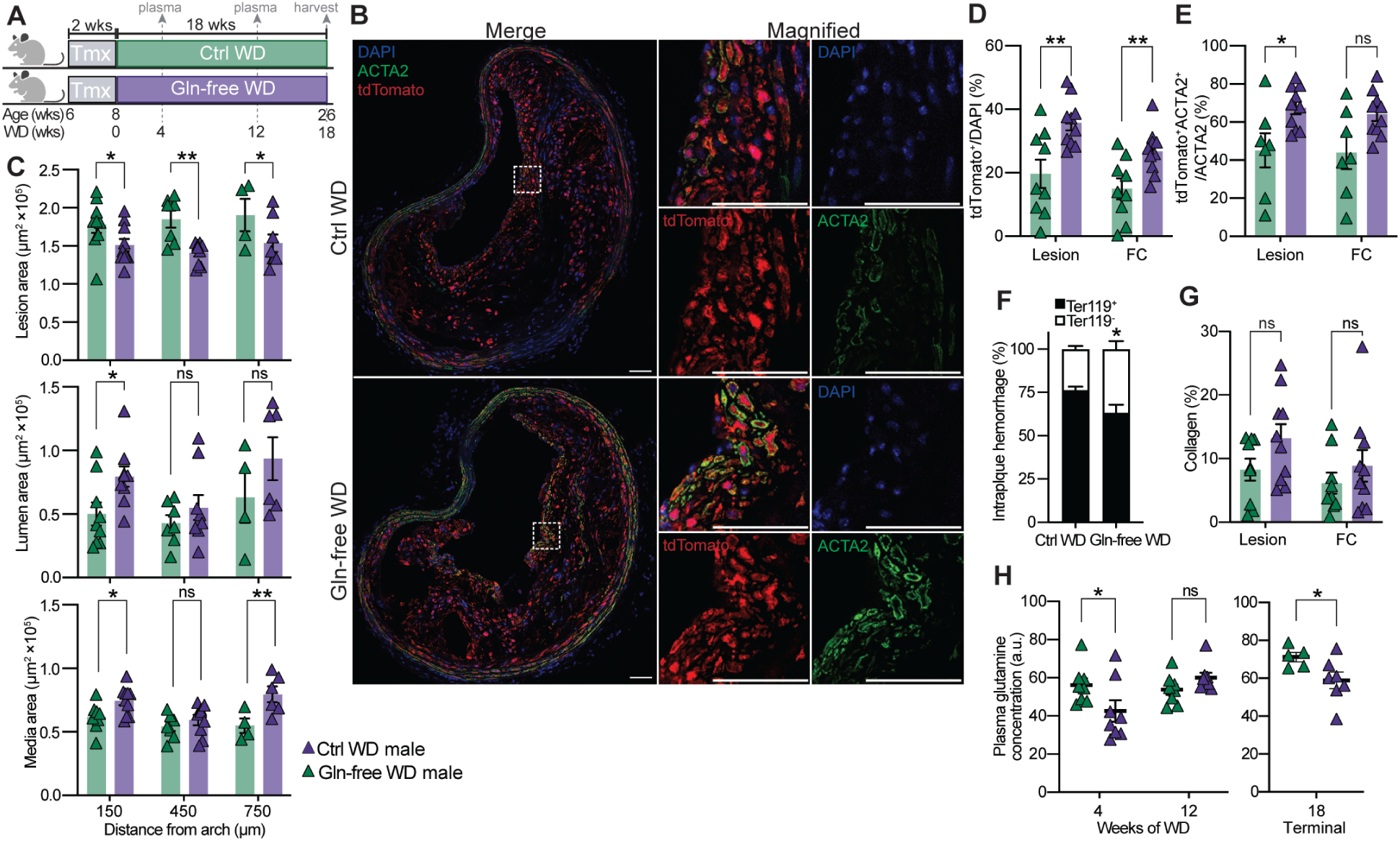
Mice fed a glutamine (Gln)-free western diet had notable increases in indices of plaque stability compared to control-diet fed mice. (**A**) Schematic of WD feeding of *Myh11*-Cre/tdTomato^+/+^/*Apoe*^-/-^ mice formulated with normal level of glutamine (2.8%; Ctrl WD) and no glutamine (Gln-free WD) for 18 weeks after tamoxifen diet to irreversibly lineage tag SMC. (**B**) Representative images of BCA lesions harvested and analyzed for SMC composition via IF staining. Regions within the white dashed square in the left images are magnified in the right images. Scale bar: 50 μm. (**C**) Vessel morphometry quantification at multiple locations distal to the aortic arch after MOVAT staining. (**D**) IF quantification of SMC and, (**E**) SMC-derived ACTA2^+^ cells in the lesion and fibrous cap. (**F**) Ter119 quantification. (**G**) No significant changes were observed for collagen deposition within the lesion and fibrous cap. (**H**) Plasma glutamine concentration collected (left) from tail vein at 4 and 12 weeks of WD, and (right) at the end of the experiment via cardiac puncture. Data analyzed using (C and F) two-way ANOVA with Tukey’s correction for post-hoc analysis or (D, E, G, H) two-tailed unpaired *t*-test. (F) Intraplaque hemorrhage was analyzed as either positive or negative staining at 2 locations past the aortic arch and *N* = 9, 8 (Ctrl WD: 150, 450 μm) or *N* = 10, 9 (Gln-free WD: 150, 450 μm). Error bars represent the mean ± SEM with *N* ≥ 5 unless otherwise indicated. \*\*\**P* < 0.001; \*\**P* = 0.001 to 0.01; \**P* = 0.01 to 0.05; ns, not significant. Graphic in (A) were created with BioRender.com.

Cell composition analysis of these lesions and fibrous cap revealed that mice fed a Gln-free WD had increased investment of SMC (*Myh11*-tdTomato*^+^*cells) and SMC-derived ACTA2^+^ cells (Fig. 4B, D to E) compared to those fed the Ctrl WD. Intraplaque hemorrhage, as assessed by Ter-119 positive staining, was reduced in Gln-free WD-fed mice compared to Ctrl WD-fed mice (Fig. 4F). Lastly, there did not seem to be a significant change in lesion collagen content in Gln-free WD-fed male mice (Fig. 4G), which is consistent with the near equal proportions of SMC-F type cells in Gln-free mice (Fig. 3H). These collective measures of plaque stability have long been used as a surrogate measure for risk of major adverse cardiovascular events. Notably, the effects of restricting dietary Gln occurred despite there being minimal reductions in plasma Gln levels in mice fed Gln-free WD for 18 weeks (Fig. 4H: right panel; data S6). Plasma Gln levels were reduced by 24% in 4-week Gln-free WD-fed mice, but by 12-weeks, the Gln levels were equivalent between the diet groups (Fig. 4H: left panel; data S7). Taken together, these results suggest that even transient decreases in dietary and plasma Gln are sufficient to change atherosclerotic plaque composition in mice.

### Atherosclerotic mice with Gln-supplemented water developed larger plaques after 8 weeks of Western diet feeding

To see whether increased dietary Gln or Gln-derived AKG would worsen lesion stability, we took *Apoe*^-/-^ mice aged 7-9 weeks and fed them a Western diet for 8 weeks while supplementing the drinking water with either 1% (w/v) AKG or 3% (w/v) Gln (fig. S8A). We found that mice with AKG containing water had 61.6% increased aortic plaque burden (fig. S8B) but no notable changes in other metrics of atherosclerosis (fig. S8, D to F). Mice with Gln water had 51.9% increased aortic plaque burden as well as 86.4% increased plaque volume within the aortic root (fig. S8, B to F). There were no detectable differences in lesion collagen (fig. S8G), necrotic area (fig. S8H) or cholesterol level (fig. S8I) across treatment groups. Given the goal is to stabilize plaques, we did not proceed with Gln supplementation for longer periods of time. These complementary outcomes of improved plaque stability with Gln depletion and decreased plaque stability with Gln supplementation predict that decreasing dietary Gln intake will decrease the rate of spontaneous plaque rupture and subsequent myocardial infarction.

### Atherosclerotic mice who experience spontaneous plaque rupture and myocardial infarction fed a Gln-free diet had improved survival compared to control-diet fed mice

Of major significance, we recently developed a murine model of atherosclerosis, major adverse cardiovascular events (MACE), and consequent mortality, termed *SR-BI*^ΔCT/ΔCT^/*Ldlr*^-/-^ ^16^. Unlike *Apoe*^-/-^ or *Ldlr*^-/-^ mice, these mice develop coronary artery lesions and exhibit spontaneous plaque rupture, myocardial infarction, and stroke after WD feeding. To test whether Gln restriction could ameliorate mortality caused by MACE, we fed these mice either Ctrl WD or Gln-free WD for up to 30 weeks (Fig. 5A). Nearly all mice fed Gln-free WD survived (13 of 14) while only 60% of Ctrl WD male mice survived the experiment (9 of 15) (Fig. 5B). Given the previously characterized cardiac toxicity in this model system, we asked whether dietary restriction of Gln results in beneficial adaptations within the heart. To address this question, we collected whole hearts from the surviving mice at the end of the experiment and performed bulk whole-tissue RNA-isolation and sequencing. The hearts from Gln-free WD fed mice had striking changes in several super-pathways, including *Metabolism* (Fig. 5C, data S8), compared to hearts from Ctrl WD-fed mice. Further dissecting changes within the *Metabolism* super-pathway, we found significant changes in the *Metabolism of Lipids*, *Carbohydrates and Carbohydrate Derivatives*, and *Amino Acids and Derivatives*(Fig. 5D, data S8). Specifically, we found that hearts from Gln-free WD-fed mice had a substantial reduction in glycosaminoglycan metabolism (forming the non-protein subunits of proteoglycans) and pyruvate dehydrogenase-dependent glucose metabolism (Fig. 5E). Additionally, these hearts had upregulated *Triglyceride biosynthesis* and pathways associated with the electron transport chain (Fig. 5, E to F). These changes in the heart transcriptome suggest that dietary Gln restriction shifts heart programming from carbohydrate metabolism to lipid metabolism. Prior studies have shown that, in the context of heart failure, hearts switch from lipid to carbohydrate catabolism, which is a less efficient form of energy production ^39,40^. In summary, removal of dietary Gln led to marked improvements in indices of plaque stability, survival, and cardiac metabolic programming in atherosclerotic mice. These results suggest that dietary reduction of Gln reduces cardiovascular mortality in mice.

**Fig 5.**
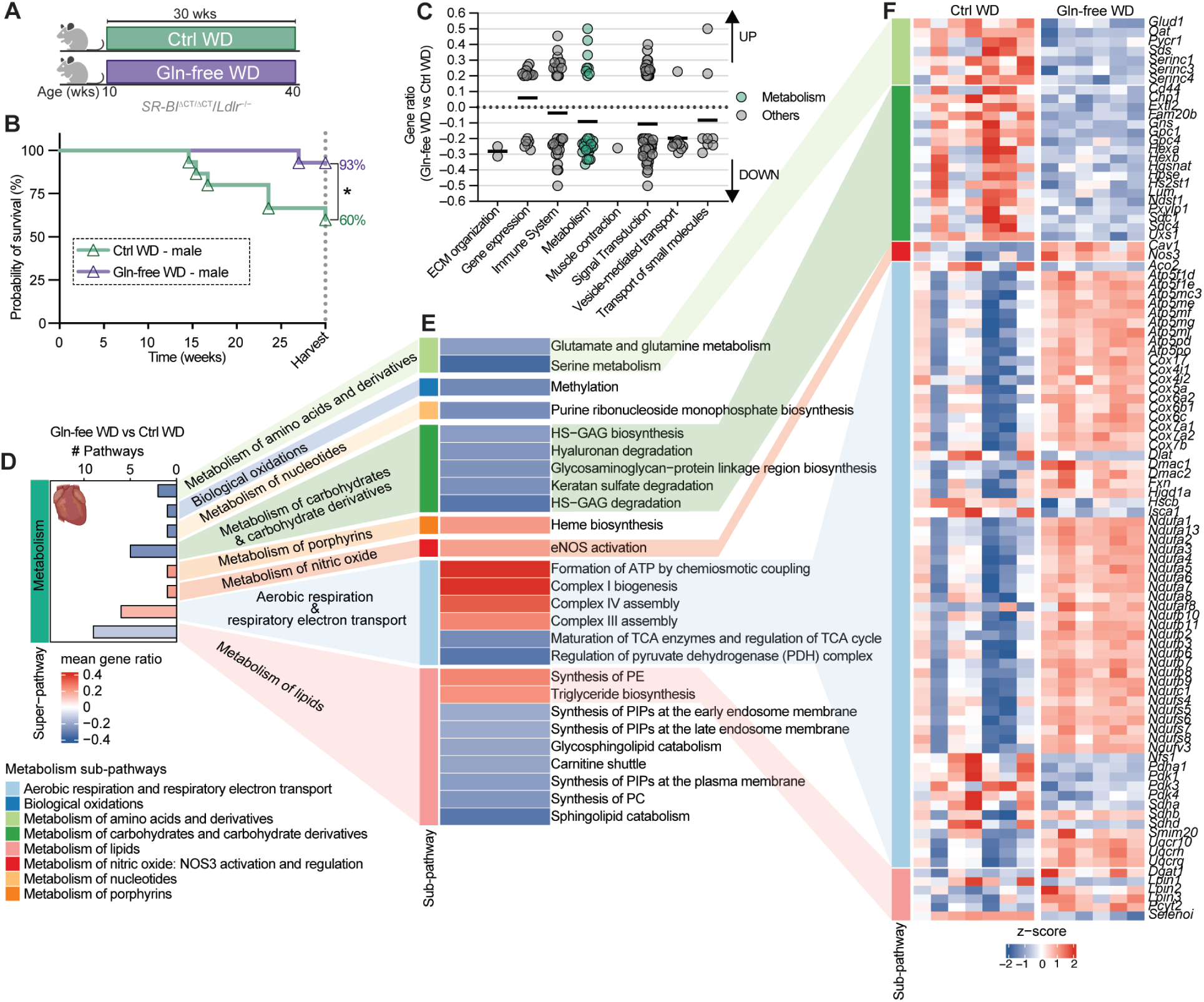
Mice fed a glutamine (Gln)-free western diet had improved survival and metabolic reprograming of the heart compared to control-diet fed mice. (**A**) Schematic of Gln-free and Ctrl WD feeding of male *SR-BI*^ΔCT/ΔCT^/*Ldlr*^−/−^ mice for 30 weeks for survival studies. (**B**) Kaplan-Meier survival curve showed that 93% of the *SR-BI*^ΔCT/ΔCT^/*Ldlr*^−/−^ male mice fed Gln-free WD survived (13/14 mice), compared with only 60% survival rate (9/15 mice) in Ctrl WD male mice. Data analyzed using (B) Kaplan-Maier curve with long-rank test. (C to F) Super-pathway analyses of bulk RNA-seq data from hearts collected from *SR-BI*^ΔCT/ΔCT^/*Ldlr*^−/−^ mice that survived 30 weeks of Gln-free or Ctrl WD. See materials and methods for super-pathway analysis and data S8 for complete results (*N* ≥ 6). (**C**) Effect of glutamine depletion on super-pathway expression pattern. Each symbol represents the gene ratio of an individual pathway within the respective super-pathway. (**D**) *Metabolism* super-pathway analysis showing the number of enriched pathways (*#pathways*) and mean gene ratio within each sub-pathway. (**E**) *Metabolism* sub-pathways. (**F**) Heatmap of differentially expressed genes (adjusted *P-*value < 0.05) grouped by selected *Metabolism* sub-pathways (complete results - data S8). Pathways in super-pathway analysis were filtered for ≥ 10 pathway size, ≥ 2 DE genes, and |gene ratio| ≥ 0.2 in at least one direction (up- or downregulated). Mean gene ratio was calculated as the average of all up- and downregulated gene ratios. \**P* = 0.01 to 0.05; ns, not significant.

### Discussion (3-5 paragraphs)

To our knowledge, this is the first study to demonstrate that dietary manipulation of a single amino acid, Gln, significantly modifies atherogenesis and survival in preclinical models. While the precise mechanisms are not fully elucidated, this finding underscores the complexity and multifaceted role of Gln in vascular pathophysiology. Gln serves not only as a nitrogen donor for the biosynthesis of nucleotides, NAD, glucosamine, and asparagine through amidotransferase activity ^41^, but also as a carbon source through its conversion first to Glu via glutaminases ^22^ and then to AKG via aminotransferases ^42,43^. To begin elucidating the complexity of Gln in atherosclerosis, we focused on SMC, which are among the most abundant cells in atherosclerotic lesions. SMC have been shown to have a pleiotropic role in atherosclerosis – serving the detrimental function of growing and destabilizing the lesion through phenotypic transitions to foam-like and pro-calcifying states and the beneficial role of stabilizing the lesion through transitions to a fibrotic state promoting ECM production ^13,14,20,44,45^.

Here, we find that Gln is critical for SMC phenotypic transitions, starting from the initial dedifferentiation (Fig. 1). This would suggest that Gln deprivation would not only restrict SMC transitions to a beneficial fibrotic state (SMC-F) but also prevent SMC-transitions to detrimental states. Further dissection of the role of Gln derived compounds found that SMC catabolize Gln-derived AKG (Fig. 2, F to K) and utilize Gln-derived Pro to synthesize collagen (fig. S5).

Notably, we find that AKG seems to play a greater role after the complete transition to SMC-F, as noted by the restoration of cell migration only after 24 hours post-treatment (Fig. 2, D and E). This is consistent with the necessity for Gln on SMC metabolic programming (Fig. 2G) and respiratory capacity after 24 hours (Fig. 2I). Considering these outcomes, it appears that AKG and Pro are Gln-derived metabolites that are required for SMC-F migration and ECM-production. However, Gln itself, possibly through other metabolites such as Glu, is critical for the dedifferentiation of SMC-C.

These findings show that that Gln provides both the substrate and the energy needed for ECM synthesis in SMC-F ^27,28^ but also permits SMC dedifferentiation and the potential to phenotypically transition to detrimental states. These in vitro studies predict that deprivation of Gln in vivo would result in a greater proportion of SMC maintaining contractile programming. Consistent with this prediction, mice fed Gln-free Western diet had a greater proportion of SMC-derived cells maintaining contractile programming with augmented migration and proliferation gene expression (migratory SMC; SMC-C2) and a smaller proportion of SMC-derived cells characterized by loss of contractile programming (dedifferentiated SMC; SMC-T) (Fig. 3H). It is possible that having more proliferative and migratory SMC can lead to a greater proportion of SMC in the lesion – as we observe in vivo – which may then adopt plaque stabilizing phenotypes. Alternatively, it is possible that inhibition of dedifferentiation of cells by lowering of Gln will lead to fewer SMC-derived detrimental phenotypes ^14,46,47^. Future studies will investigate the role of Gln in these phenotypic transitions.

The current study shows that the simple dietary strategy of Gln restriction appears to mimic the protective effects observed with hepatic *Gls2*-overexpression. Specifically, Gln absence from the diet attenuates pro-atherogenic lesion expansion and improves survival in mice. Surprisingly, we show that plasma Gln concentrations in male mice fed Gln-free diet decreased in the first 4 weeks (24%) but rebounded to the level of the Ctrl diet mice by 12 weeks (Fig. 4H; left panel). By the end of the experiment, plasma Gln levels were only minimally decreased in the Gln-free diet-fed mice compared to controls (terminal; 17.3% reduction; Fig. 4H). These latter results have several important implications. First, the initial decrease in dietary Gln triggers compensatory Gln production. Second, the long term protective effects of reduced dietary Gln are the result of stable epigenetic reprogramming of lesion cells including SMC, EC, and immune cells. Third, the beneficial effects of transient decreases in serum Gln levels on plaque stability are mediated via secondary effects in non-vascular cells, tissues, and organs. Gln is the most abundant amino acid in plasma ^48,49^ whose circulating levels are largely sustained by endogenous synthesis through Gln synthetase (*Glul*) in liver, skeletal muscle, adipose tissue, and lungs ^28^. De novo Gln synthesis accounts for 60-80% of total Gln turnover, which increases by 44-50% in healthy individuals injected acutely or chronically with cortisol to mimic moderate stress ^50,51^. These studies show that Gln demands are enhanced and often not met during hypercatabolic states including within atherosclerotic lesions ^52–55^.

Though it was not the focus of the study, our findings incidentally revealed that the effects of dietary Gln on atherosclerotic plaque stability and survival are dependent upon sex. Despite being fed a Gln-free diet or providing Gln-enriched water with WD, plasma levels of Gln were not changed in female mice (fig. S8J and fig. S9J). We assessed the indices of plaque stability in female mice, finding no major differences between the Gln-free WD and Ctrl WD groups (fig. S9, A to M; and fig. S10). Female mice also saw no survival benefit of dietary Gln restriction in the new *SR-BI*^ΔCT/ΔCT^/*Ldlr*^−/−^ mouse model that exhibits SPR, MI, and stroke (fig. S9, N and O). In summation, these studies provide novel evidence that Gln metabolism, much like SMC phenotypic switching, plays a multi-faceted role in atherosclerosis. Our deep dive into the function of Gln and Gln-derived metabolites has identified metabolic checkpoints required for SMC phenotypic switching – demonstrating the power of nuanced metabolic manipulation in controlling cell fate and function. In future studies, we will manipulate these Gln-derived metabolites to promote beneficial or inhibit detrimental SMC phenotypic transitions. Further studies are also needed to understand the role of Gln metabolism in other atherosclerosis-associated cells, such as macrophages and endothelial cells.

## Acknowledgements

This work used mass spectrometry for amino acid quantification at the Biomolecular Analysis Core (RRID: SCR_025476), cell sorting on the Becton Dickinson Influx Cell Sorter at the Flow Cytometry Core Facility (RRID: SCR_017829), library construction for scRNA-sequencing at the Genome Analysis and Technology Core (RRID: SCR_018883), and histology and immunofluorescence imaging on the Leica Thunder Imager and Zeiss LSM880 scanning confocal microscope at the Advanced Microscopy Facility (RRID: SCR_018736), all supported by the University of Virginia School of Medicine. We would like to thank Sumeet Khetarpal, Kenneth Walsh, and Clint Miller for their valuable feedback on this study.

## Funding sources

*This work was supported by:*

National Institutes of Health grants R01 HL155165 and R01 HL156849 (GKO)

American Heart Association Predoctoral Fellowship 829646 (AS)

University of Virginia Student Seed Grant-iPrime Fellowship DN001637 (AS, VM)

Swedish Research Council grant 2024-02632 (HB)

Swedish Heart Lung Foundation grant 20230337 (HB)

Royal Physiographic Society of Lund (PK)

National Institutes of Health T32 HL007284 (VM)

American Heart Association Predoctoral Fellowship 25PRE1362748 (VM)

American Heart Association Career Development Award 24CDA1269428 (VS)

## Author contribution

Conceptualization: AS, PK, HB, VMM, VS, GKO Methodology: AS, LS, HC, XY, VS

Data analysis: AS, PK, LS, VMM, NK, RR, SS, HC, DE, EB, XY, VS

Visualization: AS, PK, LS, VS Supervision: HB, SB, MM, VS, GKO

Writing – review & editing: AS, PK, HB, LS, VMM, NK, RR, SB, SS, MM, VS, GKO

Writing: AS, VS, GKO

Funding acquisition: AS, PK, HB, VMM, VS, GKO

## Data and materials availability

Transcriptomics data that support the findings of this study are available at the NCBI Gene Expression Omnibus (GEO): M437 SMC bulk RNA-seq (GSE315921); M607 SMC bulk RNA-seq (GSE316464, GSE316621); SR-BI^ΔCT/ΔCT^/Ldlr^−/−^ heart bulk RNA-seq (GSE316155); and *Myh11*-Cre/tdTomato^+/+^/*Apoe*^-/-^ BCA bulk/scRNA-seq (GSE316859, GSE318023). The mass spectrometry proteomics data have been deposited to the ProteomeXchange Consortium with identifier PXD073595.

## Extended Data for

**The PDF file includes:**

Materials and Methods

Figs. S1 to S10

Tables S1 to S2

References (*1–89*)

**Other Supplementary Materials for this manuscript include the following:**

Data S1 to S8

### Materials and Methods

#### Animal handling and tissue processing for *in vivo* studies

The metabolite supplementation in water study was approved by the regional ethics committee in Malmö/Lund (Sweden; permit number 09221/2019). The remaining animal studies were approved by the University of Virginia (UVA) Institutional Animal Care and Use Committee (USA; Animal Protocol 2400).

##### Supplementation studies

7-9 weeks old male and female *Apoe*^-/-^ mice were fed Western diet (WD) containing 21% fat and 0.2% cholesterol (Ssniff Spezialdiäten GmbH; cat#: E15721-34) for 8 weeks and received drinking water supplemented with either 3% (w/v) glutamine (Sigma; cat#: 49419), 1% (w/v) alpha-ketoglutarate (AKG) (Sigma; cat#: 75892), or control water without additives. At the end of the study, mice were euthanized by a combination of ketamine and xylazine, followed by exsanguination by cardiac puncture. The blood was collected in EDTA-containing tubes, centrifuged at 800 *g*, and plasma was stored at −80°C until further analysis. Following perfusion with PBS, hearts were stored in PBS prior to fixation, while aortas were dissected free of connective tissue and fat, opened lengthwise, and mounted *en face* for further analysis.

##### Depletion studies

Dietary glutamine depletion studies were performed on male and female mice with the following genetics: 1) C57BL/6-Tg(Myh11-cre/ERT2)F31Gko/J mice (JAX; strain #: 037658) bred to the common Rosa26-tdTomato^+/+^ and *Apoe*^-/-^ mice ^56^ are abbreviated here as *Myh11*-Cre/tdTomato^+/+^/*Apoe*^-/-^, 2) B6.FVB-Tg(Myh11-icre/ERT2)1Soff/J (JAX; strain#: 019079) bred to the common Rosa26-eYFP^+/+^ and Scarb1^tm1.1Okoch^ Ldlr^tm1Her^/J mice (JAX; strain#: 033709) ^57^, abbreviated here as *SR-BI*^ΔCT/ΔCT^/*Ldlr*^-/-^. At weaning, mice were subcutaneously implanted with micro-transponders for identification and tracking. In *Myh11*-Cre/tdTomato^+/+^/*Apoe*^-/-^ mice, Cre mediated recombination was induced by tamoxifen-containing diet (Envigo; cat#: TD.130856) provided ad libitum for 14 days starting at 6-8 weeks of age.

Following a one-week recovery period, mice were fed a custom-formulated atherogenic Western diet (WD) containing 21% fat and 0.2% cholesterol, either with (Ctrl WD, 2.8%; Envigo; cat#: TD.160725) or without glutamine (Gln-free WD, 0%; Envigo; cat#: TD.210583) for 18 weeks. These diets were designed to replicate the standard WD (Envigo; cat#: TD.88137), but with the intact protein source (casein) replaced by defined amino acids patterned after casein. The Gln-free WD contained no glutamine, with its equivalent amount (28.68 g/kg) compensated by increasing the cellulose content from 50.0 g/kg to a total of 78.68 g/kg. Male and female *SR-BI*^ΔCT/ΔCT^/*Ldlr*^−/−^ mice used for survival studies received either Gln-free WD or Ctrl WD starting at 10 weeks of age until they reached a humane terminal endpoint criteria or completed 30 weeks on the diet. Endpoint criteria for this mouse model were described previously in detail ^57^. Group assignment, data collection, and analysis were performed in a blinded manner to minimize bias. At the end of the study, *Myh11*-Cre/tdTomato^+/+^/*Apoe*^-/-^ and *SR-BI*^ΔCT/ΔCT^/*Ldlr*^−/−^ mice were euthanized via CO_2_ asphyxiation followed by exsanguination through cardiac puncture. Blood was collected in K2 EDTA-containing tubes. Collected blood was centrifuged at 300 *g* for 10 min, and stored at −80°C until analysis.

For collection of tissues for histology, we employed fixation via gravity-perfusion of the left ventricle with 5 mL of PBS, 10 mL of 4% paraformaldehyde (PFA) (EMS; cat#: 15710), and another 5 mL of PBS. Organs were harvested and weighed. BCA, and thoracis aorta, were carefully dissected and processed. Tissues were fixed for 24 hours in 4% PFA in PBS, then embedded in paraffin at the Cardiovascular Research Center (CVRC) Histology Core at UVA. **Histology**

##### BCA

4% PFA-fixed, paraffin-embedded BCAs were serially cut in 10 µm thick sections from the aortic arch to the bifurcation of the right subclavian artery. BCA sections were deparaffinized and rehydrated in xylene and ethanol series. For morphometric analysis, Modified Russell-Movat (MOVAT) staining on three locations along the BCA at 150 µm, 450 µm, and 750 µm from the aortic arch was performed at CVRC Histology Core at UVA. To assess collagen content in the lesion and fibrous cap, BCAs were stained with Picrosirius red (PSR) at CVRC Histology Core at UVA. Staining for intraplaque hemorrhage (Ter119) was performed at two locations 300 μm apart with erythroid lineage primary antibodies TER-119 (Santa-Cruz Biotechnology; cat#: sc-19592) and secondary rabbit anti-rat IgG (Vector Labs; cat#: BA-4001). Both at 1:200 dilution. Imaging of MOVAT, PSR and Ter119 was acquired on a Leica Thunder Imager at the UVA Advanced Microscopy Facility (AMF) at 10× magnification. Images were analyzed using FIJI software ^58^, as described previously ^59,60^. Briefly, images were converted to .tiff files and measurements of the lesion, external elastic lamina (EEL), and internal elastic lamina (IEL) were manually delineated on each MOVAT image using the polygon selection tool, and region of interest (ROI) was reported. The lumen area was calculated as the difference between IEL and lesion and medial thickness was calculated as the difference between EEL and IEL.

Collagen quantification was performed with custom macros on the polarized and brightfield PSR images. Predefined color threshold settings were applied to identify collagen fiber subtypes based on their birefringence properties under polarized light. Specifically, red fibers (hue 0-18, brightness 50-255), yellow fibers (hue 18-43, brightness 40-255), and green fibers (hue 43-120, brightness 40-255). Here, total collagen was reported which represents the area of positive staining for all collagen subtypes and expressed as a percentage of the total lesion or fibrous cap area. ROI for fibrous cap was determined as the 30 μm area under the lumen. Ter119 images were analyzed as Ter119 positive or Ter119 negative, and presented as percentage of staining status per location along the BCA.

##### Aortas

Oil red O was used to stain neutral lipids in en face mounted aortas. En face preparations of the aorta were fixed in 4% formaldehyde overnight, then washed in distilled water and dipped in 78% methanol. The aortas were then stained in a 0.16% (w/v) Oil Red O dissolved in 78% methanol with 0.2 mol/L NaOH.

##### Hearts

Hearts were fixed in 4% formaldehyde overnight, dehydrated and embedded in paraffin. 5 µm sections were sectioned for a total distance of 500 µm across the aortic root from the first appearance of the aortic valves. After sectioning, slides were dried at 64°C for 1 hour. To determine total plaque volume, 7 slides across the total aortic root were stained with Mayer’s haematoxylin (Histolab Products AB) and erythrosine with 3 drops of acetic acid (Acros Organics, Merck). Necrosis, defined as acellular, debris rich areas, was analysed on the slide with peak stenosis. Collagen was stained using a 2% (w/v) fast green (Sigma) solution with 2% acetic acid (Merck), followed by a 0.04% fast green, 0.1% sirius red in a saturated picric acid solution (Histolab Products AB). Slides were scanned after staining using an Aperio ScanSope digital slide scanner (ScanScope Console, Aperio Technologies, Inc.) and analysed blinded in QuPath ^61^.

#### Immunofluorescent staining

BCA sections were serially cut in 5 µm thick sections, deparaffinized and rehydrated in xylene and ethanol series. After antigen retrieval (Vector Laboratories; cat#: H-3300), sections were blocked with fish skin gelatin in 1× PBS (6 g/L) containing 10% horse serum for 1 hour at RT. Slides were then incubated overnight at 4^°^C with rabbit anti-tdTomato at 1:250 dilution (Rockland; cat#: 600-401-379). The next day the slides were washed with 1× PBS + 0.1% Tween-20 and 1× PBS for five minutes each and incubated in donkey anti-rabbit IgG Alexa Fluor 546 at 1:250 dilution, (ThermoFisher; cat#: A10040). After an additional wash step, slides were incubated in rat anti-LGALS3 at 1:500 dilution (Cedarlane; cat#: CL8942AP) for one hour. Donkey anti-goat conjugated to Alexa 647 at 1:250 dilution (Invitrogen; cat#: A21447), mouse monoclonal SM α-actin-FITC at 1:250 dilution (ACTA2) (clone 1A4, Sigma Aldrich; cat#: F3337), and DAPI at 1:100 dilution (5 mg/mL; Invitrogen; cat#: D21490) were all added after a wash step and left to incubate for one hour at RT. After final wash, slides were mounted using Prolong Gold Antifade (Invitrogen; cat#: P36934). Slides were imaged within one week on a Zeiss LSM880 scanning confocal microscope at the University of Virginia AMF core with a 20× objection at a resolution of 2048×2048 16 bit depth, 11 sections with a step of 1 µm.

##### Cellular composition analysis

Immunofluorescent images were converted in .tiff files and lesion ROIs were generated using FIJI software. The images and ROIs were then run through a convolutional neural network trained from datasets in the Owens laboratory as described previously ^62,63^ to determine the tdTomato positive SMC contribution to the atherosclerotic plaque. Data was then loaded into the Colocalization Object Counter plugin ^64^ for FIJI where ACTA2^+^ cells were labeled by hand.

#### Plasma analysis

For supplementation studies, glutamine was analyzed using Glutamine Assay Kit, according to the manufacturer’s instructions (Abcam; cat#: ab197011). Plasma cholesterol was analyzed, using Infinity Cholesterol Liquid Stable Reagent kit (Thermo Scientific; cat#: TR13421) and Data-Trol A, Abnormal Control Serum (Thermo Scientific; cat#: TR41001), as an internal standard, according to the manufacturer’s instructions. For depletion studies, lipids and liver enzymes quantification were performed at the UVA Clinical Pathology Laboratory using an automated hematology analyzer (Sysmex; cat#: XN-9000). Amino acids levels were quantified using targeted PRM (Parallel Reaction Monitoring) mass spectrometry (MS) at the Biomolecular Analysis Facility.

##### Targeted Amino Acid Quantification via PRM-MS

Plasma samples (10 µL) were extracted using a modified biphasic methanol-chloroform protocol. Briefly, 80 µL of cold 100% methanol was added to each sample, followed by vortexing and incubation at 4 °C for 15 minutes with shaking at 900 rpm. Subsequently, 40 µL of cold chloroform and 80 µL of water were added, and samples were vortexed and centrifuged at 10,000 rpm for 10 minutes to induce phase separation. The resulting upper aqueous phase was dried in SpeedVac. Dried extracts were reconstituted in 20 µL of 0.1% formic acid (2× the initial volume) containing 10 µM of a heavy-labeled amino acid mixture (Cambridge Isotope Laboratories; cat#: MSK-CAA-1) which served as internal standards for quantification. A 5 µL aliquot of each reconstituted sample was injected for analysis.

Chromatographic separation was achieved using a Waters ACQUITY UPLC BEH C18 column. The mobile phases consisted of 0.1% formic acid in water (Solvent A) and 0.1% formic acid in methanol (Solvent B). The column was maintained at 30°C, and separation was performed at a flow rate of 0.25 mL/min with a 9 minute gradient of 5-98% B.

Full MS scan spectra were acquired at 60,000 resolution over an m/z range of 67-1000, with an RF lens setting of 35%, AGC target of 50%, and maximum injection time of 50 ms in positive ion mode. PRM scans were acquired at 30,000 resolution using a collision energy of 30 ± 10, 1.6 m/z isolation window, RF lens of 35, AGC target of 50%, and 50 ms maximum injection time.

Processing of raw files was done in Skyline software v23.1 ^65^. The concentration of each amino acid was calculated based on the ratio of its peak area to that of its corresponding heavy internal standard. The ratio was multiplied by the concentration of the internal standard (10 µM). A final multiplication factor of 2 was applied to correct for the twofold sample concentration during the reconstitution step. Cysteine, glycine, and alanine were excluded from the final quantification due to poor retention/peak shape. Absolute concentrations were compiled and reported in data S6-7.

#### In vitro

##### Cell Culture

SMC, referred here as M607, were isolated from *Myh11*-eYFP^+/+^/*Apoe*^+/+^ mice, as previously described ^59^. Briefly, thoracic aortas from 20 mice were dissected, and the adventitia was digested using an enzyme cocktail containing 1 mg/mL Collagenase-2 (Worthington Biochemical Corp; cat# 4202), 0.75 U/mL Elastase ES (Worthington Biochemical Corp; cat#: 2279), and 1 mg/mL Soybean Trypsin Inhibitor (Worthington Biochemical Corp; cat#: 3571). After digestion, the adventitia was removed, vessel was cut open *en face* and the endothelial layer on inside of aorta was gently scraped away using forceps. Aorta was washed three times with PBS, then digested further to a single-cell suspension.

Cells were initially plated and cultured in DF20 [DMEM/F12 media (Gibco; cat#:11320033) supplemented with 20% fetal bovine serum (FBS; HyClone; cat#: SH30071.03)], followed by maintenance of cells in DF10 [DMEM/F12 with 10% FBS]. Both DF20 and DF10 were further supplemented with 2 mM L-glutamine (Gibco; cat#: 25030081) and 100 U/mL of pen/strep (Gibco; cat#:15140122).

To ensure pure SMC-population, cells were assessed by the Invitrogen Countess 3 FL Automated Cell Counter equipped with an EVOS YFP light cube (Invitrogen; cat#: AMEP4954) and showed >90% eYFP^+^ SMC.

New SMC cell line, referred here as M437, was isolated as above. M437 was isolated from one thoracis aortas of male *Myh11*-tdTomato^+/+^/*Apoe*^-/-^ mouse and pure SMC population was assessed with an EVOS RFP light cube (Invitrogen; cat#: AMEP4952).

##### Treatments

SMC were plated on tissue culture plates and grown in DF10 to reach a confluence before being switched to serum free media (SFM) for 24-72 hours. SFM was prepared by supplementing DMEM/F12 with: 2 mM L-glutamine, 100 U/mL pen/strep, 0.2 mM L-ascorbic acid (Sigma; cat# A4403), 5 µg/mL transferrin (Sigma; cat# T5391), 16 µg/mL insulin (Gibco; cat# 12585014), and 6.25 ng/mL sodium selenite (Sigma; cat# S5261). Cells were washed with PBS and treated with one or a combination of the following, depending on the experiment: PDGF-BB (30-50 ng/mL; vehicle – 0.5% BSA in 4 mM HCl/PBS), TGFB (10-50 ng/mL; vehicle – 0.5% BSA in 4 mM HCl/PBS), glutamine (2 mM; vehicle – water), DMKG (2 mM; vehicle – DMSO), BPTES (5 µM; vehicle – DMSO), CB-839 (1 µM; vehicle – 0.5% BSA in 4 mM HCl/PBS), NFLP (50 mM; vehicle – PBS), AOA (1 mM; vehicle – water), or vehicle control. For more details about products used for cell treatments see table S1.

##### Quantitative Real-Time PCR

Approximately 1×10^5^ SMCs were lysed in RLT buffer (Qiagen; cat#: 74136), and RNA was isolated according to the manufacturer’s protocol. RNA was reverse-transcribed using iScript cDNA Synthesis Kit (Bio-Rad; cat#: 1708891). qRT-PCR was performed using SensiFast SYBR No-ROX (Bioline; cat#: BIO-98050) and run on a Bio-Rad CFX56 Real-Time System. The analysis was performed using 2^−ΔΔCt^ method with beta-2-microglobulin (*B2m*) for normalization. Primer sequences for qPCR as well as genotyping are listed in table S2.

##### Glycolytic and Mitochondrial Stress Test

Approximately 4×10^4^ or 1×10^4^ SMCs were seeded onto a Seahorse XF24 (Agilent Technologies XFe96/XF Pro FluxPak; cat# 103792) or XFe96 well cell culture (Agilent Technologies XFe24 FluxPak mini; cat# 102342) in DF10 and grown to confluence. SMC were serum starved for at least 24 hours before treatments for 0-72 hours. Respiratory and glycolytic capacities were assessed using DMEM+Gln media (20 mM glucose; 2 mM glutamine) or DMEM-Gln media (20 mM glucose; 0 mM glutamine; Thermo-Fisher cat#: 12800017, pH = 7.35 ± 0.02 at 37°C) using a Seahorse XF24 and XF96 Flux Analyzer (Agilent Technologies; Santa Clara, CA) as established previously ^59^ and briefly described here.

The Mitochondrial Stress Test (MST) measured the oxygen consumption rate (OCR) of cells, representing mitochondrial respiratory ability. OCR was measured initially (basal respiration), and after injection of 1 μM oligomycin (Sigma-Aldrich; cat#: 75351), 2 μM BAM15 (respiratory capacity; Cayman Chemical Company; cat#: 17811), as well as after injection of 10 μM antimycin A (Sigma-Aldrich; cat#: A8674) & 1 μM rotenone (non-mitochondrial; used for normalization, Sigma-Aldrich; cat#: R88751G). The contribution of glutamine to respiratory capacity was determined by subtracting the X + BPTES (Sigma-Aldrich; cat#: SML0601) condition’s respiratory capacity from the X + vehicle condition, where X is vehicle, or PDGF/TGFB. The contribution of glucose was determined as the respiratory capacity of the X + BPTES condition. Glycolytic stress test (GST) measuring extracellular acidification rate (ECAR) of treated SMC, represent glycolytic ability. ECAR was measured initially in the absence of glucose, after injection of 20 mM D-glucose (basal glycolysis), 1 μM oligomycin (glycolytic capacity), and 80 mM 2-deoxy-D-glucose (non-glycolytic; used for normalization; Sigma; cat#: D8375). Energy capacity map was used to represent the bioenergetic potential of SMCs using glycolytic capacity (ECAR) and respiratory capacity (maximal OCR).

##### Quantification of secretome from conditioned cell media

Approximately 450 μL, of cell culture media post-treatment was collected for analysis of secreted proteins. Samples were concentrated down to 35 μL using Amicon Ultra Centrifugal Filters (3 kDa MWCO, Millipore Sigma; cat#: 239UFC003). Bradford protein quantification assay was employed to quantify protein levels.

Briefly, concentrates were subjected to reduction and alkylation using 10 mM DTT and 50 mM iodoacetamide, and a tryptic digestion followed by C18 clean-up using Bravo AssayMAP C18 cartridges (Agilent) for peptide purification ^66^. Samples were separated by reversed-phase HPLC (U3000 RSLCnano, 75 μm x 50 cm EASY-Spray C18 RP column) and subjected to a field asymmetric ion mobility spectrometry (FAIMS, Thermo Scientific) for gas-phase fractionation followed by data-independent acquisition (DIA) on a Thermo Scientific Orbitrap Fusion Lumos Tribrid Mass Spectrometer. Raw data were analyzed using Spectronaut (v18, Biognosys) and protein abundances were exported for further analysis. Protein abundances output from Spectronaut were used for quantification. To avoid potential misidentification of FBS from tissue culture, bovine proteins were removed and mouse serum proteins with bovine homologs were also removed if: 1) the bovine homolog was detected in the majority of samples while the mouse homolog was missing; 2) the bovine homolog showed >10-fold higher abundance than the mouse homolog. Analysis was performed in R (v.4.4.2). Statistical tests for each comparison were performed independently. Protein with quantified values in > 3 samples in at least one condition were subjected to log_2_-transformation, normalization, and imputation of missing values. Normalization was performed using the method of variance stabilizing transformation from vsn package ^67^. Imputation was performed on proteins present in one condition while absent in another condition (off-condition) by randomly imputing one sample in the “off-condition” with a background-level signal using deterministic minimum method (ImputeLCMD package) ^68,69^. Principal component analysis (PCA) was performed to evaluate global variations and batch effects prior to imputation. Differential expression analysis was performed using limma package ^70^. *P*-values from the empirical Bayes moderated *t* test were adjusted for multiple-testing with the Benjamini-Hochberg method (BH). Proteins with adjusted *P*-values < 0.05 were considered statistically significant.

#### Live cell imaging and analysis

##### Cell Migration Assay

Approximately 1×10^4^ SMCs were seeded onto an Imagelock 96-well cell culture plate (Sartorius; cat#: BA-04856) in DF10 and grown to confluence. M607 and/or M437 cells were serum starved for at least 24 hours before treatments for 0-72 hours. Assay was performed using Incucyte® 96-well Woundmaker Tool which creates 96 cell-free zones in cell monolayers. Cells were then washed with PBS and treated with 100 µL of SFM containing experimental treatments. Cells were imaged at every 2-3 hours for up to 4 days in Incucyte® S3 Live-Cell Analysis System (Sartorius; Göttingen, Germany) using phase contrast imaging as well as green fluorescence channels for M607 SMC or red for M437 at 10× objective. The Incucyte ® Scratch Wound analysis module was then used for real-time, automated measurement of wound confluence ^71^.

##### Cell eccentricity quantification

The base analysis software from Incucyte® S3 Live-Cell Analysis System was also used to quantify cells morphology metric - eccentricity (cells roundness) on the phase and fluorescent images captured during the migration assay at 10× objective.

##### Cell morphology classification

Approximately 8×10^4^ SMCs were seeded onto a 24-well cell culture plate (Corning) in DF10 to reach confluence. Cells were then serum-starved for ≥ 24 hours before administration of treatments. Phase-contrast images were acquired every 2-3 hours across 72 hours using an Incucyte® S3 live-cell imager with a 10× objective; 2-4 non-overlapping images were captured per well. For analysis, the Incucyte® Cell-by-Cell analysis software module was used which applies a segmentation mask to identify individual cells per image. To distinguish cell morphologies, we applied the Incucyte® Advanced Label-Free Classification module. Classifier employs multivariate analysis to identify multiple morphological features of cells such as symmetry, texture, area, brightness, perimeter length, and circularity, which are used to train a classifier through control wells ^72,73^. The classifier was trained on a set of control images: vehicle treated contractile SMC representing class A (SMC-C; circular morphology) and PDGF/TGFB treated cells as class B (SMC-F; non-circular, spindle-like morphology) (fig. S2A). Classifier was then applied to all wells of treatment groups and timepoints. The model computes a classification score between 0 and 1 for each cell per image, and a threshold of 0.3 was empirically established to distinguish class B (SMC-F; non-circular cells) from class A (SMC-C). The proportion of class B cells per image across replicates was calculated and is presented as the percentage of F-like cells (% SMC-F).

#### RNA isolation for RNA-sequencing

BCA samples for RNA extraction were collected from male and female *Myh11*-Cre/tdTomato^+/+^/*Apoe*^-/-^ mice fed either Gln-free WD or Ctrl WD for 18 weeks. Heart tissue for RNA extraction was collected from male *SR-BI*^ΔCT/ΔCT^/*Ldlr*^-/-^ mice fed either Gln-free WD or Ctrl WD for 30 weeks. Mice were perfused solely with PBS (20 mL), and tissues were snap-frozen in liquid nitrogen and stored at −80°C.

RNA from BCA tissue was homogenized using TissueLyser II (Qiagen) at 30 Hz for 6 minutes and RNA was extracted using RNeasy Micro Kit (Qiagen; cat#; 74004). To isolate RNA from cells, RNeasy Plus Mini Kit was used (Qiagen; cat#: 74136). All steps were performed according to the manufacturer’s protocol. RNA extraction from frozen heart tissue was performed by Novogene using TRIzol-based high-lipid protocol.

#### RNA-seq analysis

Sequencing was conducted by Novogene. Samples that passed the RNA quality check were used for polyA-enriched mRNA library preparation, followed by sequencing on Illumina NovaSeq 6000, except for RNA-seq from M437 SMC samples and frozen heart tissue, which were sequenced on NovaSeq X-Plus platform. Sequencing specifications included paired-end reads at 150bp length (Illumina PE150 technology) and data output of at least 25 million reads pair per sample. Fastq files were filtered using fastp v0.23 ^74^ to remove 1) uncertain nucleotides (*N*) that constitute more than 10 percent of either read (*N* > 10%), 2) low quality nucleotides (phred quality score < 20), and 3) reads shorter than 15bp. Quality of the reads was assessed using FastQC v0.11 ^75^ and MultiQC v1.14 ^76^ software. Filtered reads were aligned to the mouse mm39 reference genome (genome assembly GRCm39) using STAR v2.7 software ^77^, and counted using featureCounts from Subread package v2.0 ^78^. Downstream analyses were performed in R Studio v4.3 ^79^. Differentially expressed genes (DEGs) were identified by DESeq2 package v1.42 ^80^ after removing low count genes, defined as counts with expression lower than 10 in a minimal number of samples per group. Wald-test *P*-values were adjusted for multiple tests using the Benjamini-Hochberg (BH) correction, and genes with a 5% false discovery rate (adjusted *P*-value < 0.05) were considered significant. For visualization, counts were normalized using variance stabilizing transformation (VST) and standardized to z-scores for heatmap generation.

#### Pathway and Super-Pathway Analysis

Sets of genes for pathway analysis were obtained from the Molecular Signatures Database v2024.1.Mm ^81^ and included two curated sources: the Reactome pathway database (12,981 total gene sets) and the Matrisome collection of ECM (9 total gene sets), the latter of which includes 938 unique EMC-genes not present in the ECM collection from the Reactome database.

##### Gene Set Enrichment Analysis (GSEA)

GSEA was performed using the GSEA function from the clusterProfiler v4.10 package ^82^. Prior to GSEA, genes were filtered to include only those with a baseMean expression ≥ 15. The DESeq2 Wald statistic values were used to rank genes. GSEA was performed using the default parameters including BH correction, except for the following modifications: minimum gene set size was set to 14, and eps was set to 0. The resulting enriched gene sets were reported as Normalized Enrichment Score (NES) and were considered significantly enriched when adj *P*-value < 0.05. For each significantly enriched gene set, the gene ratio was calculated from the core enrichment genes (also known as the “leading-edge subset”) divided by all annotated genes within each pathway.

##### Super-Pathway Analysis

To summarize differentially expressed genes (DE; adjusted *P*-value < 0.05) from bulk RNA-seq and scRNA-seq data, we developed and performed a custom “Super-Pathway” framework. This analysis groups enriched pathways based on Reactome’s hierarchical structure where top-level pathways are composed of multiple smaller pathways, which in turn may be subdivided into even smaller pathways. Each enriched pathway was traced through up to 11 hierarchical levels to identify its top-level parent (termed “super-pathway”) and immediate child (termed “sub-pathway” or “subcategory”). Each sub-pathway may itself be a parent pathway to another set of sub-pathways. Each pathway, irrespective of level, has a finite non-zero number of annotated genes – the number of these annotated genes within a pathway is referred to here as pathway size. We represent the above concepts mathematically in the relationships below, where *X* is any pathway and *Y* is any sub-pathway belonging to *X*:

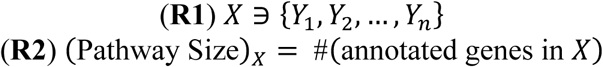

In practice, this means that multiple specific pathways enriched in our analysis could be summarized under a single super-pathway. Pathway hierarchy relationships were obtained from the Reactome database v91 ^83^. To integrate Matrisome pathways into the analytical framework, we defined analogous groupings based on established ECM classifications in Matrisome Project^84^. For example, “Naba_Proteoglycans” were grouped under the subpathway “Naba_Core Matrisome” and under the super-pathway “Naba_Matrisome”.

Using the abbreviation of GR for “gene ratio”, for each terminal pathway (not as a parent or group of pathways), we calculated the following metrics:

1. the upregulated gene ratio (“GR-UP”), which represents the proportion of upregulated genes and is defined by **Equation 1**.

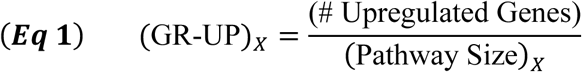
2. the downregulated gene ratio (“GR-DOWN”), which represents the proportion of downregulated genes and is defined by **Equation 2**.

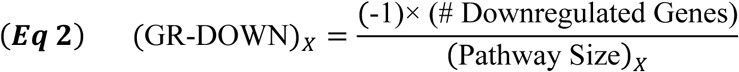
3. net gene ratio (referred to as “gene ratio” in the text), represents overall pathway regulation and was defined as a sum of gene ratio up and gene ratio down as shown in **Equation 3**.

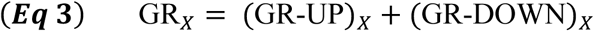

Finally, we calculated mean gene ratio (MGR) for any super-pathway *X* or non-terminal sub-pathway, which represents the summation of the net gene ratio for each sub-pathway *Y* divided by the number of sub-pathways belonging to *X*.

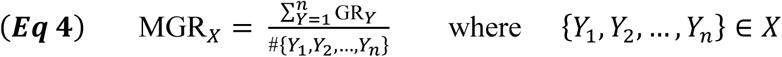

The pathways are considered to be significantly changed if: 1) minimum pathway size ≥10 genes (gene set), 2) minimum of 2 genes within pathways were significantly changed (adjusted *P*-value < 0.05 based on DESeq2 analysis), and 3) pathways with GR-UP ≥ 0.2 or |GR-DOWN| ≥ 0.2 (i.e. ≥20% of genes within this pathway were significantly changed; adjusted *P*-value < 0.05). Values not meeting these filtering criteria were set to NA. This Super-Pathway analysis framework was applied to bulk RNA-seq data performed on M437 and M607 SMC cells as well as to scRNA-seq data to compare SMC phenotypes found in murine atherosclerotic lesions.

#### scRNA-sequencing

##### Cell processing

*Myh11*-Cre/tdTomato^+/+^/*Apoe*^-/-^ male and female mice were fed Gln-free WD and Ctrl WD for 18 weeks (as shown in Fig. 4A; *N* = 4 mice per group) and harvested to isolate a single cell suspension from individual BCA lesion. BCA lesion were collected and placed into FACS buffer (1% BSA in PBS) on ice until all samples were collected. Tissues were chopped with scissors and dissociated in a digestions cocktail containing 4 units/mL Liberase (Roche; cat#: 355374) and 0.744 units/mL Elastase (Worthington Bio. Corp; cat#: 2279) for 60 min at 37°C. FACS buffer was added and each sample was filtered through a 40 μm filter insert followed by centrifugation for 10 min at 500*g*. Pallet was resuspended and incubated in RBC lysis buffer (BD Biosciences; cat#: 555899) for 5 min at RT. FACS buffer was added, followed by centrifugation for 10 min at 500 *g*. Each sample was resuspended in Cell Multiplexing Oligo (CMO) from CellPlex Kit Set A (10x Genomics; cat#: 1000261) and incubated for 5 min at RT. FACS buffer was added, followed by centrifugation for 10 min at 500*g*. Next, each sample was resuspended in PBS, viability dye was added (Invitrogen; cat#: 65086514) and incubated for 10 min at RT in dark. Each sample was washed with PBS and centrifuged at 500*g* for 10 min.

Individually labelled samples were resuspended in FACS buffer and pooled to four experimental groups: Ctrl WD males, Gln-free WD males, Ctrl WD females, Gln-free WD females. Each group had 4 samples assigned with oligonucleotide tag CMO309-CMO312. Live cells were sorted using a Becton Dickinson Influx Cell Sorter, counted and submitted directly for scRNA-seq library preparation.

##### Library construction

Gene Expression (GEX) and Cell Multiplexing Libraries (CMO) were prepared using the Chromium Next GEM Single Cell 3ʹ Reagent Kits v3.1 (Dual Index; TT Set A index for GEX, NN Set A index for CMO) with Feature Barcode technology for Cell Multiplexing (protocol CG000388 Rev C) at the Genome Analysis and Technology Core at the UVA, in accordance with the manufacturer’s instructions.

##### Sequencing

We aimed to capture 18,000 cells per group (4,500 cells per sample), targeting approximately 100,000 reads per cell for GEX and 10,000 reads per cell for CMO libraries. Libraries were pooled and run equally across 7 lanes on the NovaSeq X Plus sequencing system (Illumina) at Novogene with system specifications of paired-end reads at 150bp length (PE150) and a total data output of 2598 Gb.

##### Read alignment

Fastq files were aligned to a custom reference mouse genome (GRCm39) with the reporter gene tdTomato-WPRE-polyA using Cell Ranger v8.0. *Myh11*-Cre/tdTomato^+/+^/*Apoe*^-/-^ mice were generated by crossing with a tdTomato reporter from mouse known as Ai14 (The Jackson Laboratory, strain#: 007914) ^56^. tdTomato-WPRE-polyA sequence was obtained from the Ai9 vector (Addgene; cat#: 22799) since the Ai9 mouse shares the same sequence for tdTomato-WPRE-polyA with the Ai14 mouse used in this study ^85^.

Fastq files were demultiplexed using the cellranger multi pipeline (default parameters) to identify each sample by their oligonucleotide tag (CMO). Percentage of cells assigned to any of the four CMOs per group was low: 46% for Ctrl F Group, 0% for Ctrl M Group, 0.01% for Gln-free F Group, 57% for Gln-free M group. Because we pooled samples based on their experimental groups and constructed them as separate libraries, we proceeded with cellranger count pipeline instead of cellranger multi (v8.0) and we detected > 13,000 cells per group.

##### scRNA-seq data processing

Raw count matrices were imported to R v4.3 utilizing Seurat package v5.1. Seurat object included all the libraries with additional two healthy control libraries collected previously in our lab from BCA of chow fed female and male mice. Initial filtering removed genes that do not occur in a minimum of 5 cells. Then, cells with fewer than 400 detected genes, and 500 UMIs were removed. The maximum threshold was set as median+3*MAD (median absolute deviation). We also applied the threshold for percent of mitochondrial genes as maximum of 15% and 5% for hemoglobin genes. We used the DoubletFinder ^86^ to remove doublets using an estimated multiple rate of 8% as expected multiplet rate is 0.4% for every 1,000 targeted cells. Further analytical processing included normalization with SCTransform v2 ^87^ with 10,000 variable features, and integration using the reciprocal PCA (rpca) workflow. Eighteen clusters were initially identified using FindNeighbors and FindClusters functions with resolution = 0.3, and principal components (PCs) = 35 from a total of 98,602 cells. Cluster 16 was the smallest (271 cells), and cluster 8 was the biggest in terms of number of cells (18,858 cells). To analyze cells of SMC origin, we excluded tdTomato^-^cells, and re-clustered the remaining 9,642 cells at 0.3 resolution and 22 PCs. It was previously reported that Rosa26 locus in the tdTomato reporter line has a baseline leak, and cells with log-normalized tdTomato levels above 3.5 were defined as tdTomato^+^ cells ^88^. In our SCTransform-normalized data, the equivalent threshold would be 3.04. However, we applied a more conservative threshold of 3.18, corresponding to the 90th percentile of tdTomato expression, to ensure specificity. This approach identified 5 tdTomato^+^ clusters. Clusters were identified based on canonical markers from literature-curated and statistically ranked genes. Differential expression analysis was performed using FindAllMarkers function with Model-based Analysis of Single-cell Transcriptomics (MAST) test on genes expressed by more than 50% of cells in either of the two populations (min.pct = 0.5), and log_2_ fold change ≥ 0.25. Pathway and super-pathway analysis was performed as stated above. We performed pseudotime analysis on the UMAP embeddings using Monocle 3 v1.3.7 ^89^, specifying cells of SMC1 cluster as roots of the pseudotime analysis.

#### Statistical Analysis and Reproducibility

Statistics were performed using GraphPad Prism software or R Studio. The sample size is represented by the number of data points within individual graphs or is indicated in the figure legend. Data points represented with a square symbol indicate results from female mice; data points represented as triangles represent male mice. For cell culture results, data points represented by circles symbols indicate results generated from the M607 cell line, whereas diamond symbol was used for results generated from the M437 cell line. Each in vivo murine and in vitro cell culture experiment was repeated at least twice. Data are presented as mean ± SEM or median ± IQR (fig. S8). The sample size and type of statistical analysis performed are reported within each figure. *P*-value ≤ 0.05 was considered statistically significant.

**Fig. S1.**
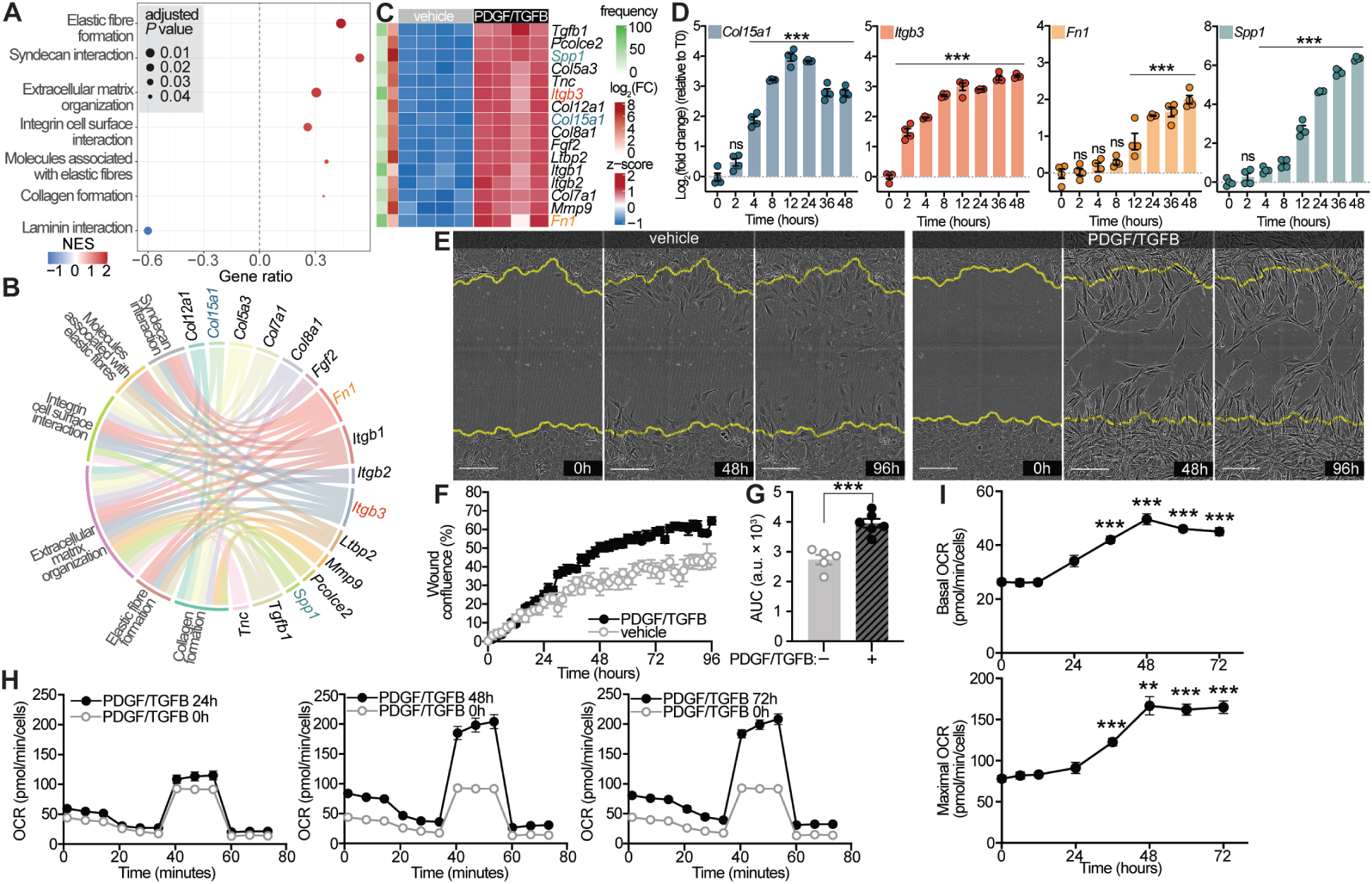
Aortic SMC, when treated with PDGF/TGFB are characterized by enhanced expression of ECM genes, as well as increased respiratory and migratory capacities, related to Figure 1. (A, B, C) Bulk RNA-seq analyses after PDGF/TGFB treatment at 24 hours in M607 SMC. (**A**) Dot plot displaying pathways enriched in Gene Set Enrichment Analysis (GSEA) related to extracellular matrix (ECM). NES: Normalized Enrichment Score. (**B**) ECM marker-pathway network displaying marker genes that participate in multiple ECM-related pathways. (**C**) Heatmap showing the expression of identified ECM marker genes. Row annotations display: (1) pathway frequency - the percentage of ECM pathways in which each gene appears as a leading edge gene, and (2) log_2_(FC) of PDGF/TGFB versus vehicle at 24 hours; FC is fold change. In (B) and (C) only marker genes that met the criteria of appearing in > 50% of ECM related pathways (pathway frequency) or having > 25% pathway frequency with |log_2_(FC)| > 3 are included. Highlighted genes correspond to the selected markers in (D) and are color-coded consistently. (**D**) Time-course expression of selected ECM-markers after PDGF/TGFB treatment plotted as log_2_(FC) relative to PDGF/TGFB treatment at 0-hour (T0). (**E**) Representative images of M607 from migration assay at 0, 48, and 96 hours after PDGF/TGFB treatment. The yellow line indicates the initial borders of the scratched area. Scale bar: 200 μm. (**F**) Change in wound confluence over time was measured, and (**G**) area under the curve was calculated after 96 hours of treatment with PDGF/TGFB, a.u.: arbitrary unit. (**H**) Mitochondrial stress test measuring oxygen consumption rate (OCR) at 12, 24, 48, and 72 hours post PDGF/TGFB treatment. (**I**) Basal respiration (initial OCR measurements), and (bottom) respiratory capacity, measured after injection of oligomycin, BAM15, and antimycin A & rotenone (non-mitochondrial; used for normalization) of PDGF/TGFB treated M607 at various time points relative to 0-hour. Data analyzed using (G) two-tailed unpaired *t*-test or (I) one-way ANOVA with Dunnett’s correction for post-hoc analysis with *N* ≥ 4. Error bars represent the mean ± SEM. \*\*\**P* < 0.001; \*\**P* = 0.001 to 0.01; \**P* = 0.01 to 0.05; ns, not significant.

**Fig S2.**
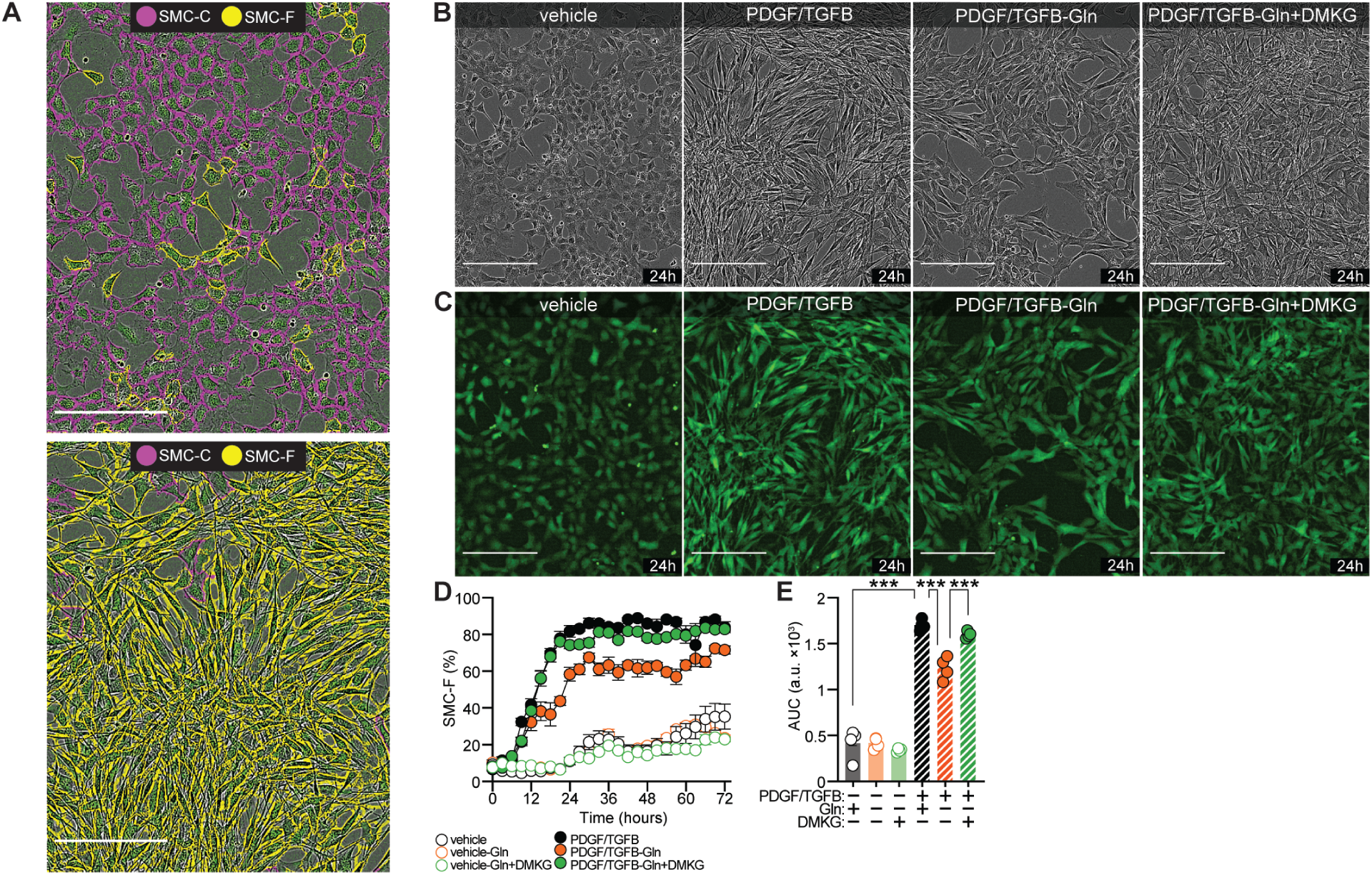
Morphology-based classification of PDGF/TGFB-induced SMC phenotypic transitions, related to Figure 1. (A to F) M607 SMC were treated with or without PDGF and TGFB, Gln, or DMKG. Images were acquired every 3 hours for 3 days. (**A**) Incucyte® advanced label-free classification was performed to identify and quantify different cell morphologies of two different SMC phenotypes: circular SMC-C (pink) and non-circular, spindle-like SMC-F (yellow). Scale bar, 200 μm. (**B** and **C**) Representative images showing (B) phase contrast and (C) eYFP fluorescence, taken after 24 hours of treatment. Scale bar, 200 μm. (**D**) Quantification of the percent of SMC-F cells during the 3 day treatment. (**E**) Area under the curve (AUC) analysis of (D), a.u.: arbitrary unit. Data analyzed using one-way ANOVA with Tukey’s correction for post-hoc analysis with *N* ≥ 4, error bars represent mean ± SEM. \*\*\**P* < 0.001.

**Fig. S3.**
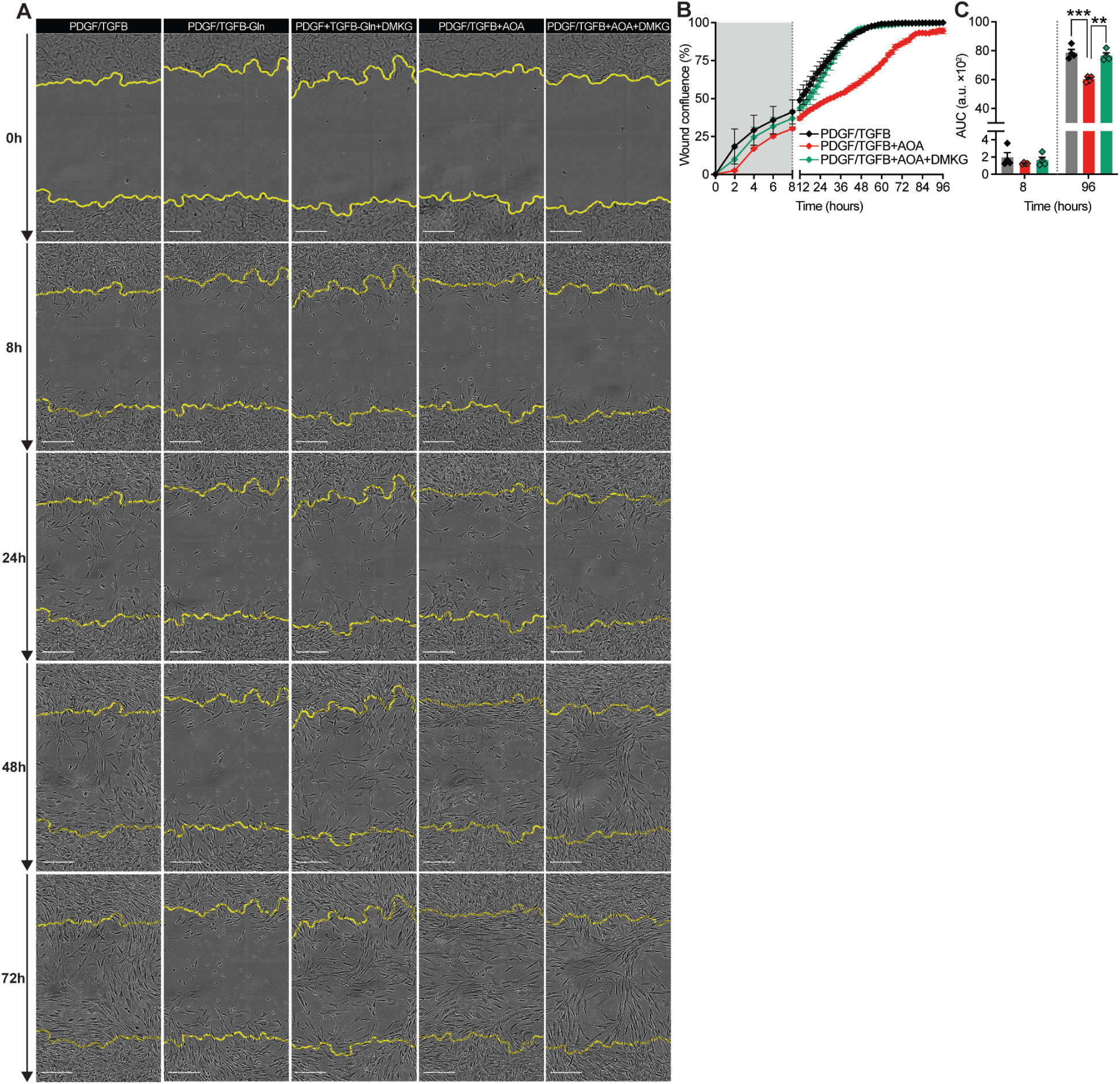
DMKG rescues the migration deficiency in SMC treated with PDGF/TGFB in the absence of Gln, related to Figure 2. (**A**) Representative phase contrast images of M437 SMC treated with PDGF/TGFB in the presence or absence of glutamine (-Gln), AOA, and DMKG. Images shown at 0, 8, 24, 48, 72 hours post-wounding in a scratch wound assay. Initial wound edge marked by yellow line. Scale bar = 200 μm. (**B**) Wound closure kinetics over time in the presence of AOA and AOA+DMKG in PDGF/TGFB treated SMC. The grey shading highlights the early response that is Gln-independent. (**C**) Area under the curve (AUC) analysis of scratch closure after 8 and 96 hours for treatment conditions shown in panels (A) and (B), a.u.: arbitrary unit. Graphs were analyzed using one-way ANOVA with Tukey’s correction for post-hoc analysis. Error bars represent mean ± SEM with *N* ≥ 4. \*\*\**P* < 0.001; \*\**P =* 0.001 to 0.01; \**P =* 0.01 to 0.05.

**Fig. S4.**
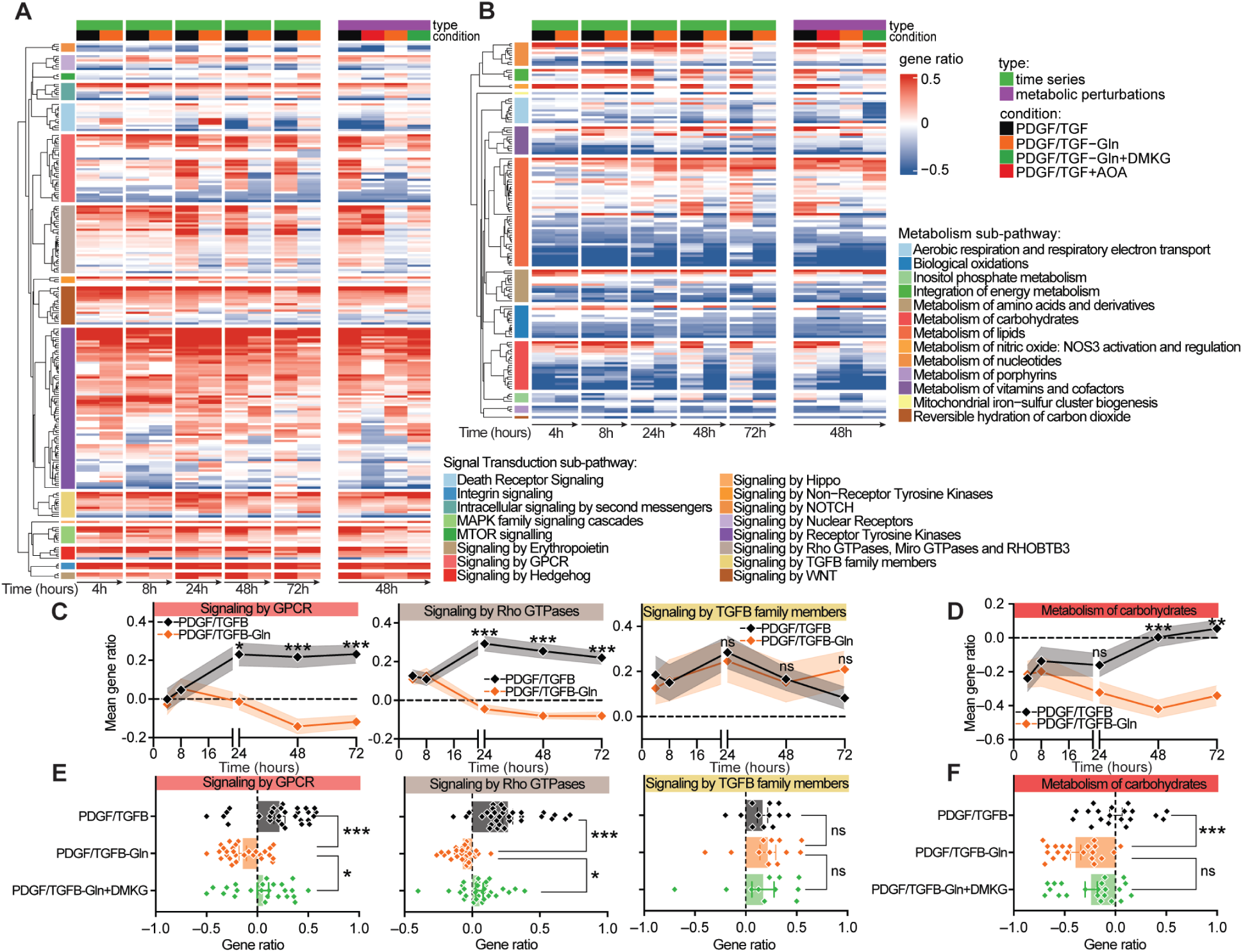
Analysis of dynamic transcriptional *Signal Transduction* and *Metabolism* sub-pathways during SMC-F transition following manipulation of glutamine metabolism in M437, related to Figure 2. (A to F) Super-pathway analyses of bulk RNA-seq data from M437 treated with PDGF/TGFB in the presence or absence of glutamine, DMKG, or AOA. See materials and methods for super-pathway analysis and data S3-S4 for complete results. (**A**) Heatmap of *Signal Transduction* and (**B**) *Metabolism* super-pathways, organized by all sub-pathways across time (left), and after metabolic perturbations with AOA and DMKG at 48 hours (right). (**C**) Selected *Signal Transduction*, and (**D**) *Metabolism* sub-pathways. Mean gene ratio was calculated as the average of all pathway gene ratios within each sub-pathway. (**E** and **F**) Bar plots showing the rescue efficiency of DMKG for each sub-pathway from (C) and (D). Each diamond symbol represents the gene ratio of an individual pathway within the respective sub-pathway. Rows in (A) and (B) represent individual pathways hierarchically clustered within sub-and super-pathways. Gene ratios represent the proportion of significantly changed genes relative to all annotated genes within each pathway. Positive gene ratio (red) indicate net upregulation and negative values (blue) indicate net downregulation relative to 0 hour. Pathways in super-pathway analysis were filtered for pathway size ≥10 genes, ≥ 2 DE genes, and |gene ratio| ≥ 0.2 in at least one direction (up- or downregulated). Data analyzed using (C and D) two-way ANOVA with Sidak’s or (E and F) one-way ANOVA with Tukey’s correction for post-hoc analysis. Error bars represent mean ± SEM. \*\*\**P* < 0.001; \*\**P =* 0.001 to 0.01; \**P* = 0.01 to 0.05; ns, not significant.

**Fig. S5.**
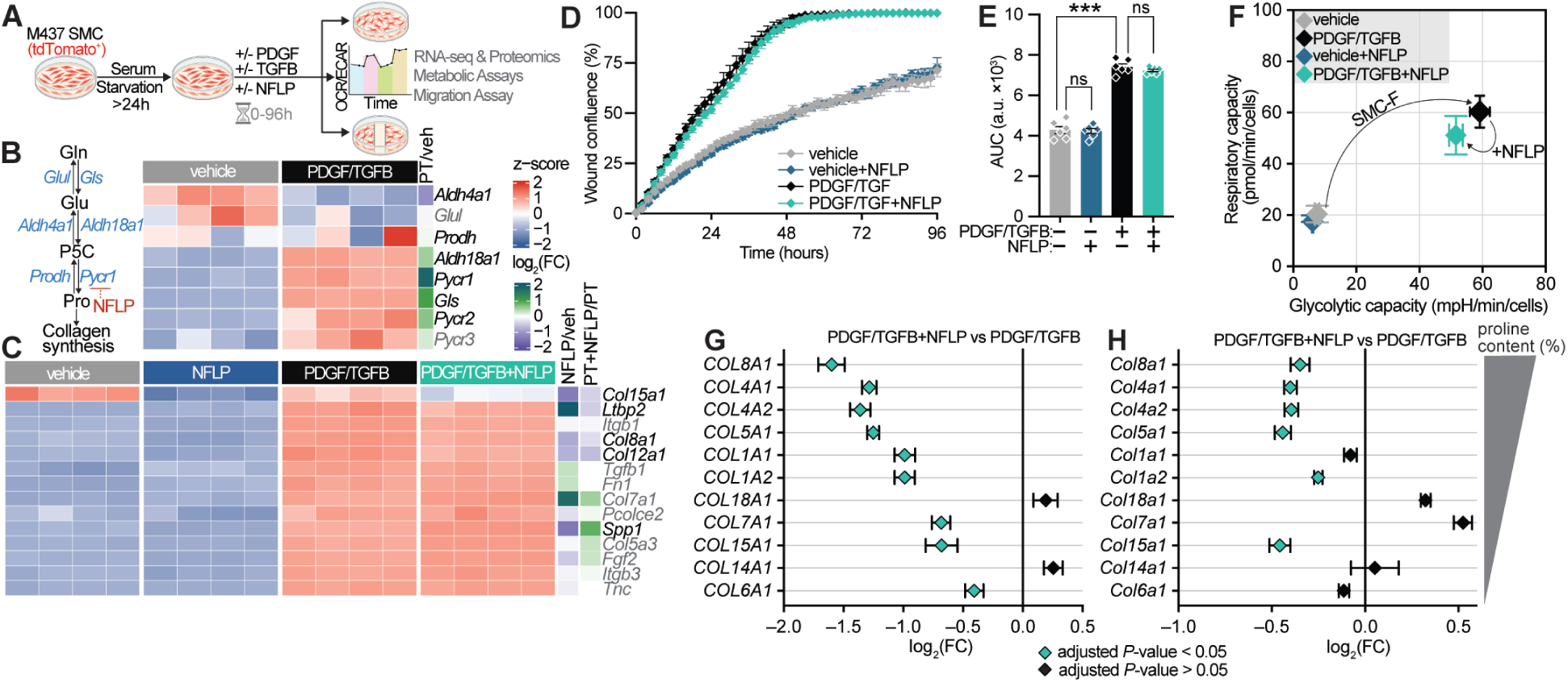
The Gln-Glu-Pro axis is essential for certain collagens to be produced by SMC, but not for the overall transition to SMC-F state, related to Figure 2. (**A**) Experimental design. SMC (line M437; derived from aorta of *Myh11*-Cre/tdTomato^+/+^/*Apoe*^-/-^ male mice) were treated with PDGF/TGFB in the presence or absence of N-Formyl-L-proline (NFLP) which inhibits glutamate-to-proline (Glu-Pro) conversion. After 48 hours, bulk RNA-seq, proteomics, and bioenergetics analyses were performed; migration assay was assessed up to 96 hours after treatment. (B and C) Bulk RNA-seq analysis of genes involved in (**B**) Gln-Glu-Pro synthesis and utilization. Pyrroline 5-carboxylate (P5C) is the intermediate product of both anabolism and catabolism of proline (Pro). P5C is synthesized from glutamate by aldehyde dehydrogenase 18 family, member A1 (*Aldh18a1*; also knows as pyrroline-5-carboxylate synthase) and then converted to proline (Pro) by pyrroline-5-carboxylate reductase (*Pycr1, Pycr2, Pycr3*). Pro is degraded by pyrroline-5-carboxylate dehydrogenase (*Prodh*) to P5C and to glutamate (Glu) via aldehyde dehydrogenase 4 family, member A1 (*Aldh4a1*; also knows as pyrroline-5-carboxylate dehydrogenase). (**C**) Expression of EMC-markers after 48 hours of treatment with NFLP. In (B) and (C), genes in grey denote genes of interest that were not statistically significantly changed (adjusted *P*-value > 0.05). Row annotations display: (1) z-score, and (2) log_2_(FC) of PDGF/TGFB versus vehicle (PT/veh), NFLP versus vehicle (NFLP/veh), and PDGF/TGFB+NFLP versus PDGF/TGFB (PT+NFLP/PT). FC: fold change. (**D**) Change in wound confluence over time was measured, and (**E**) area under the curve was calculated after 96 hours of treatment. (**F**) Energy capacity map representing the bioenergetic potential of SMC after 48 hours, using glycolytic capacity (x-axis; stressed ECAR) and respiratory capacity (y-axis; maximal OCR). (**G** and **H**) Forest plot ordered by the collagens with the highest to lowest proline content and representing the effect of NFLP in SMC-F at the (G) protein, and (H) gene level. Collagens with the significant effect were highlighted in teal shade (adjusted *P*-value < 0.05). Data analyzed using one-way ANOVA with Tukey’s correction for post-hoc analysis with *N* ≥ 4, error bars represent mean ± SEM. \*\*\**P* < 0.001; \*\**P* = 0.001 to 0.01; \**P* = 0.01 to 0.05; ns, not significant.

**Fig. S6.**
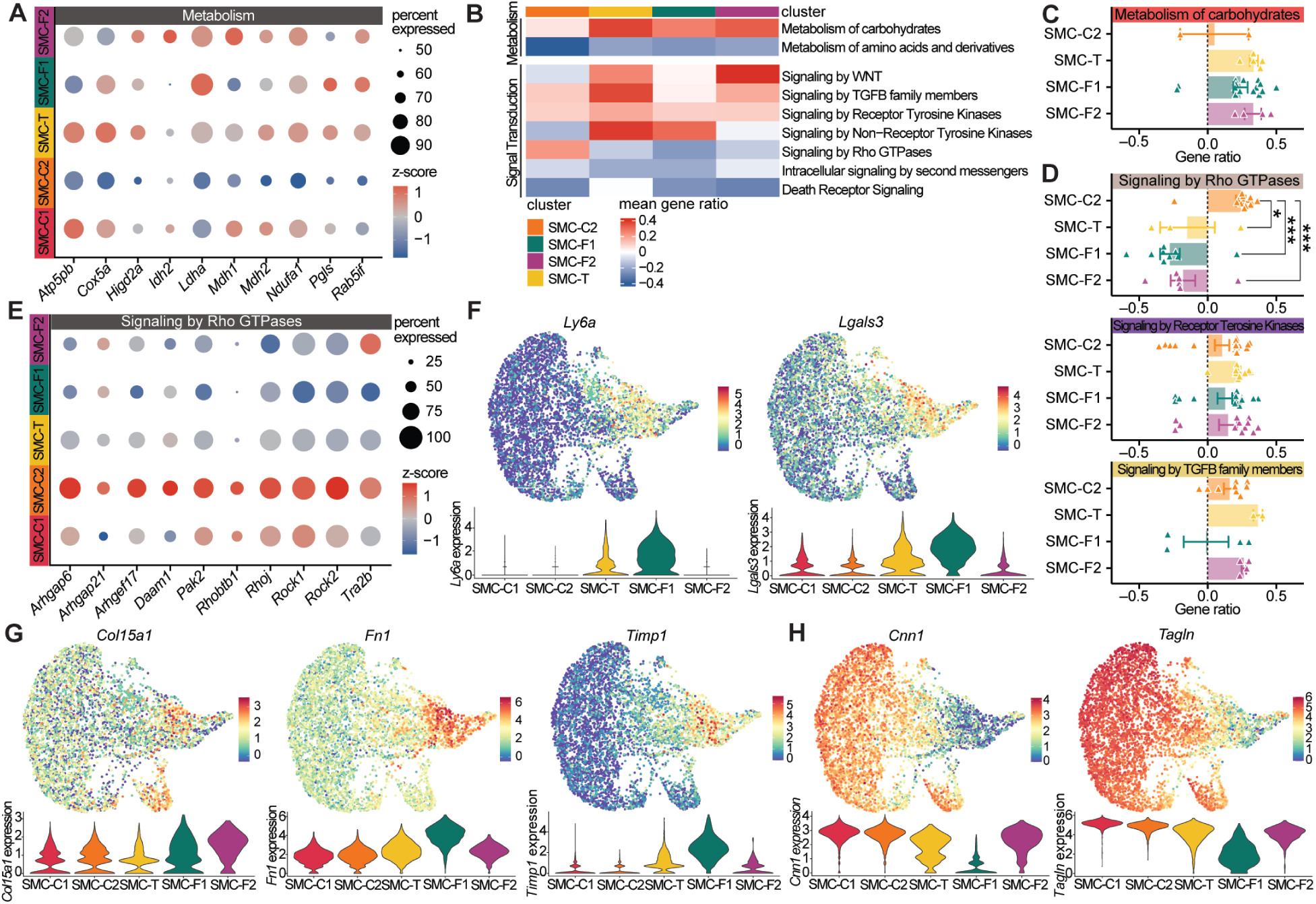
scRNAseq analysis of SMC-derived clusters (tdTomato^+^) present within atherosclerotic brachiocephalic arteries of male mice, related to Figure 3. (**A**) Dot plot showing scaled gene expression related to metabolism in SMC-clusters. (**B**) Heatmap of sub-pathways within *Metabolism* and *Signal Transduction* super-pathways across SMC clusters. Only sub-pathways with more than one pathway enriched in all clusters shown. Positive mean gene ratio (red) indicate net upregulation and negative values (blue) indicate net downregulation. (**C**) Bar plots of *Metabolism of carbohydrates*, and (**D**) *Rho GTPases-, Receptor Tyrosine Kinase-,* as well as *TGFB family member signaling* correspond to the selected sub-pathways from panel B. Each symbol represents the gene ratio of an individual pathway within the respective sub-pathway. (**E**) Dot plot showing scaled gene expression related to *Signaling by Rho GTPases.* (**F**) Feature and violin plots of SCTranformed gene expression of *Ly6a* and *Lgals3,* as well as (**G**) ECM-markers, and (**H**) contractile markers. (A to D) Data derived from differential expression analysis focusing on genes expressed by more than 50% of cells and with a log_2_(FC) ≥ 0.25 in 3 male samples. SMC-C1 was used as the baseline reference point for comparing each cluster (SMC-C2, SMC-T, SMC-F1, and SMC-F2). (B and C) Super-pathway analyses of SMC clusters from scRNA-seq data. Pathways were filtered for pathway size ≥10 genes, ≥ 2 DE genes, and |gene ratio| ≥ 0.2 in at least one direction (up- or downregulated). See materials and methods for super-pathway analysis and data S5 for complete results. In (C) and (D) data analyzed using one-way ANOVA with Tukey’s correction for post-hoc analysis. Error bars represent mean ± SEM. \*\*\**P* < 0.001; \*\**P =* 0.001 to 0.01; \**P* = 0.01 to 0.05; ns, not significant.

**Fig. S7.**
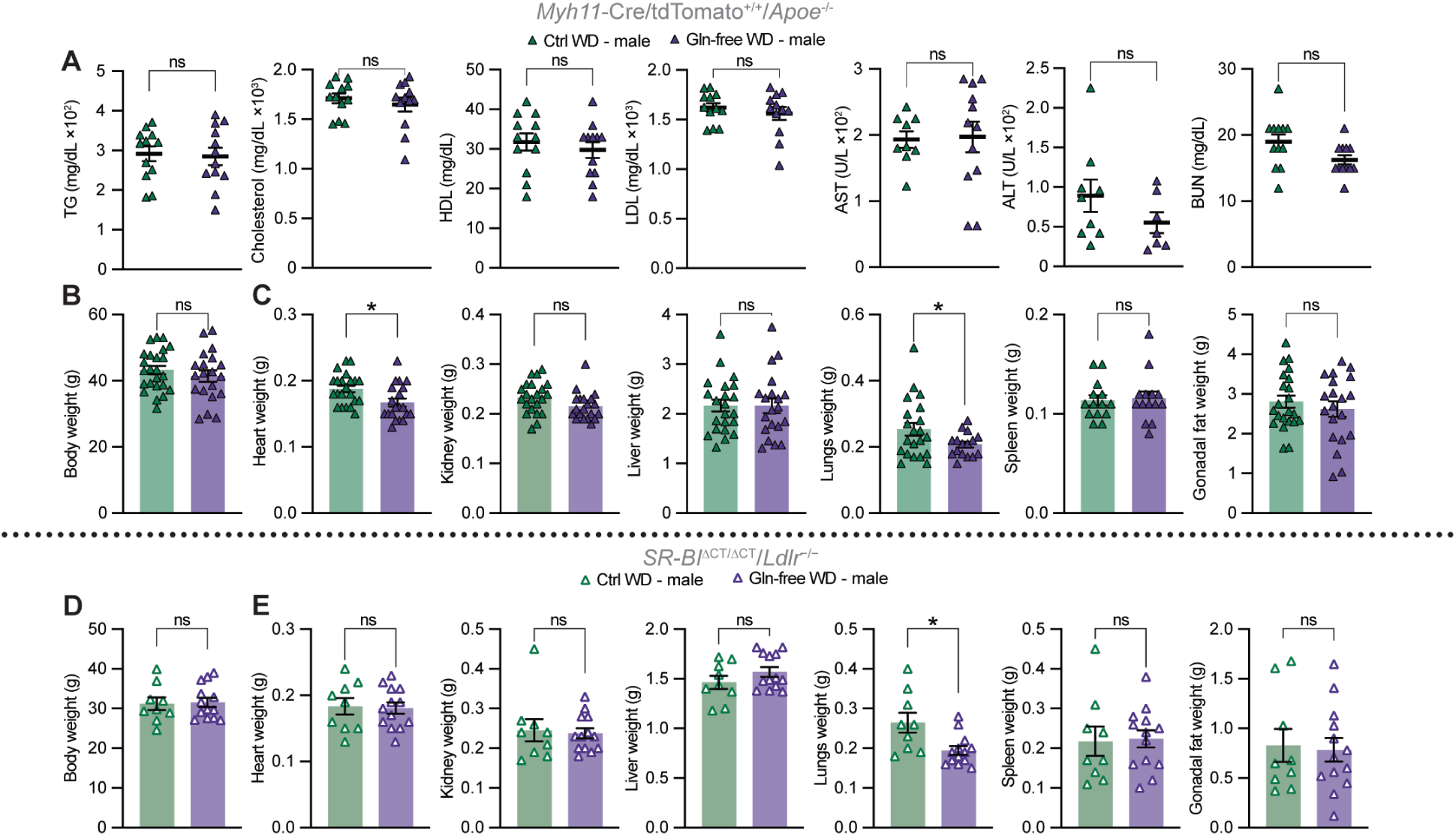
Plasma metabolic panel and mouse parameters in male mice fed a Western diet (WD) with and without glutamine, related to Figure 4 and 5. (A to H) After 18 weeks of WD, *Myh11*-Cre/tdTomato^+/+^/*Apoe*^-/-^ male mice showed (**A**) no significant changes in plasma markers between experimental groups (TG: Triglycerides, HDL: High Density-, and LDL: Low-Density Lipoprotein, AST: Aspartate Transaminase, ALT: Alanine Transaminase, BUN: Blood Urea Nitrogen; *N* ≥ 7 per group). (**B**) Depletion of glutamine had no effect on body weight after 18 weeks of WD (*N* ≥ 20 per group). (**C**) Organ weights, including kidney, liver, spleen, and gonadal fat, were unaffected by Gln-free WD, but weights of heart and lungs were significantly reduced compared to Ctrl WD. *N* ≥ 14 per group. (**D**) After 30 weeks of WD, *SR-BI*^ΔCT/ΔCT^/*Ldlr*^−/−^ male mice showed no effect on body weight between experimental groups. (**E**) Organ weights were unaffected except for reduced lung weight (*N* ≥ 9 per group; mice surviving the experiment from Fig. 5B). Data analyzed using two-tailed unpaired *t*-test. Error bars represent the mean ± SEM. \**P =* 0.01 to 0.05; ns, not significant.

**Fig. S8.**
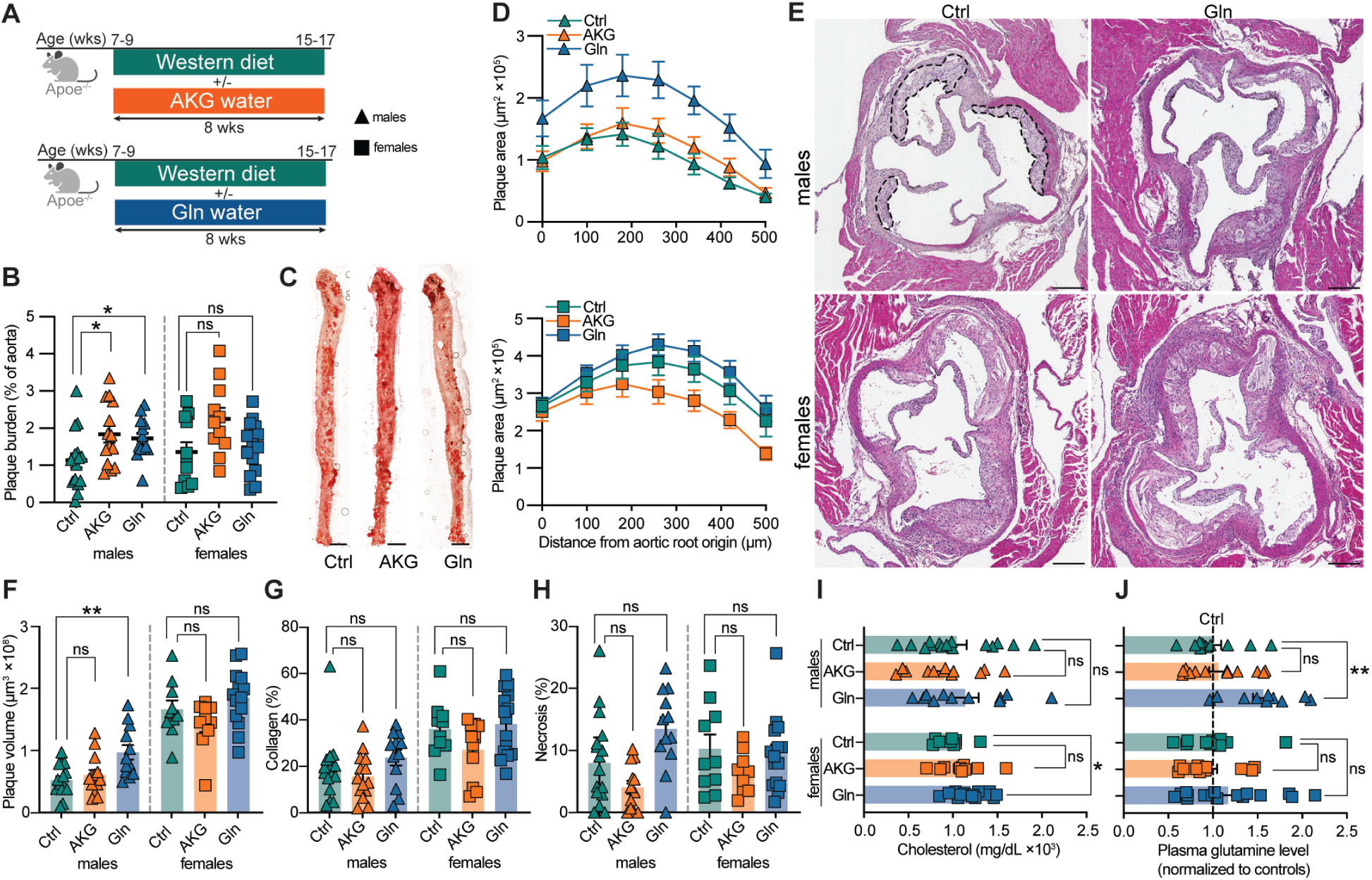
Atherosclerotic mice with supplemented glutamine in water developed larger plaques after 8 weeks of Western diet feeding, related to Figure 4. (**A**) Experimental design. *Apoe*^-/-^ male mice were fed WD and received drinking water with no additives (Ctrl) or water supplemented with glutamine (Gln), or alpha-ketoglutarate (AKG) for 8 weeks. (**B**) Oil red O analysis of plaque burden in the aorta with (**C**) representative images of aortas from male mice. Scale bar: 2 mm. (**D**) Plaque area at 7 locations across the aortic root from the first appearance of the aortic valves in (top) male and (bottom) female mice. (**E**) Representative images of aortic root plaques from male and female mice in control (Ctrl) and glutamine (Gln) group. Scale bar: 200 μm. Black dashed line marks representative plaques boundaries. (**F**) Total plaque volume in the aortic root, calculated as the area under the curve from (D). (**G**) Sirius red analysis of plaque collagen content. (**H**) Necrotic core size analysis at the point of peak stenosis. (**I**) Plasma cholesterol. (**J**) Glutamine plasma concentration normalized to sex matched controls. (B, F, G, I, J) Data analyzed using one-way ANOVA with Dunnett’s correction or (H) Kruskal Wallis with Dunn’s correction for post-hoc analysis with *N* ≥ 13. Error bars represent mean ± SEM or (H) median ± IQR. \*\*\**P* < 0.001; \*\**P* = 0.001 to 0.01; \**P* = 0.01 to 0.05; ns, not significant.

**Fig. S9.**
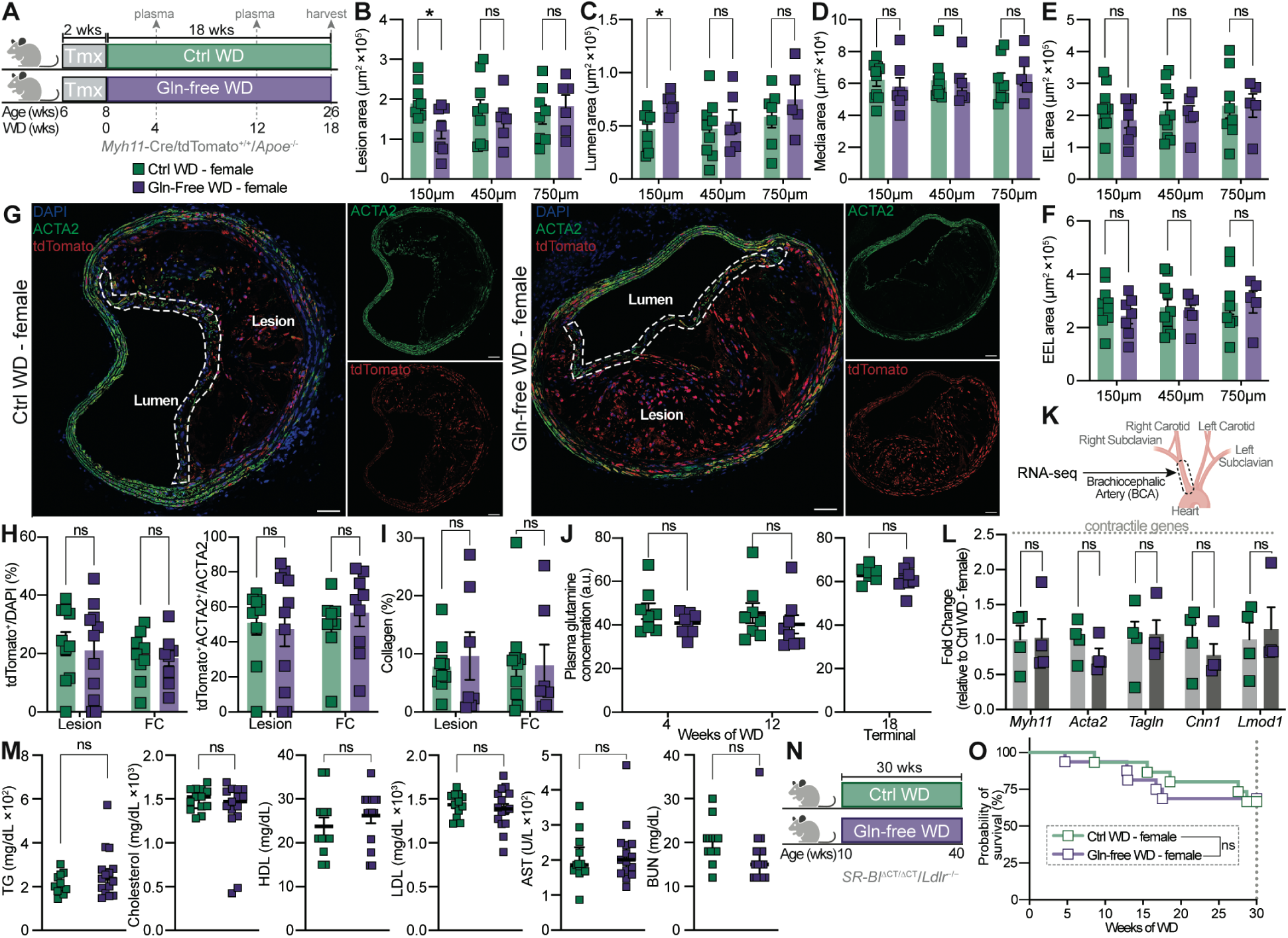
Female atherosclerotic mice fed a glutamine (Gln)-free Western diet showed no effect on indices of plaque stability and survival, related to Figure 4. (**A**) Schematic of Ctrl WD and Gln-free WD feeding of *Myh11*-Cre/tdTomato^+/+^/*Apoe*^-/-^ female mice for 18 weeks. (B to F) Vessel morphometry: (**B**) lesion, (**C**) lumen, (**D**) media thickness, (**E**) internal-, and (**F**) external elastic lamina at 3 locations past the aortic arch. (**G**) IF representative images of BCA. White dashed line marks fibrous cap. Scale bar: 50 μm. (**H**) IF quantification of (left) SMC and, (right) SMC-derived ACTA2^+^ cells in the lesion and fibrous cap (FC). (**I**) Collagen deposition. (**J**) Plasma Gln concentration collected (left) from tail vein at 4 and 12 weeks of WD, and (right) at the end of the experiment via cardiac puncture (a.u.: arbitrary unit). (**K**) Schematic of BCA dissection for bulk RNA-seq after 18 weeks of WD, and (**L**) expression levels of contractile genes presented as fold change relative to Ctrl WD. *N* = 4. DESeq2 results showed no statistical changes: *Myh11* log_2_(FC) = 0.031, *P* = 0.97; *Acta2* log_2_(FC) = -0.39, *P* = 0.3; *Tagln* log_2_(FC) = 0.11, *P* = 0.89; *Cnn1* log_2_(FC) = -0.37, *P* = 0.53; *Lmod1* log_2_(FC) = 0.2, *P* = 0.77. FC: fold change. (**M**) Plasma metabolic panel after 18 weeks of WD. TG: Triglycerides, HDL: High Density-, and LDL: Low-Density Lipoprotein, AST: Aspartate Transferase, BUN: Blood Urea Nitrogen. (**N**) Survival study schematic: *SR-BI*^ΔCT/ΔCT^/*Ldlr*^−/−^ female mice fed Ctrl WD or Gln-free WD for up to 30 weeks. (**O**) Kaplan-Meier curve showed no effect on survival, specifically 69% of Gln-free WD female mice (11/16 mice) survived, compared with 67% survival rate in Ctrl WD female mice (10/15 mice). (B to F) Data analyzed using two-way ANOVA with Tukey’s correction, (H, I, J, M) unpaired *t*-test, (O) log-rank test. Error bar represent the mean ± SEM, *N* ≥ 7 unless noted. \**P* < 0.05; ns, not significant.

**Fig. S10.**
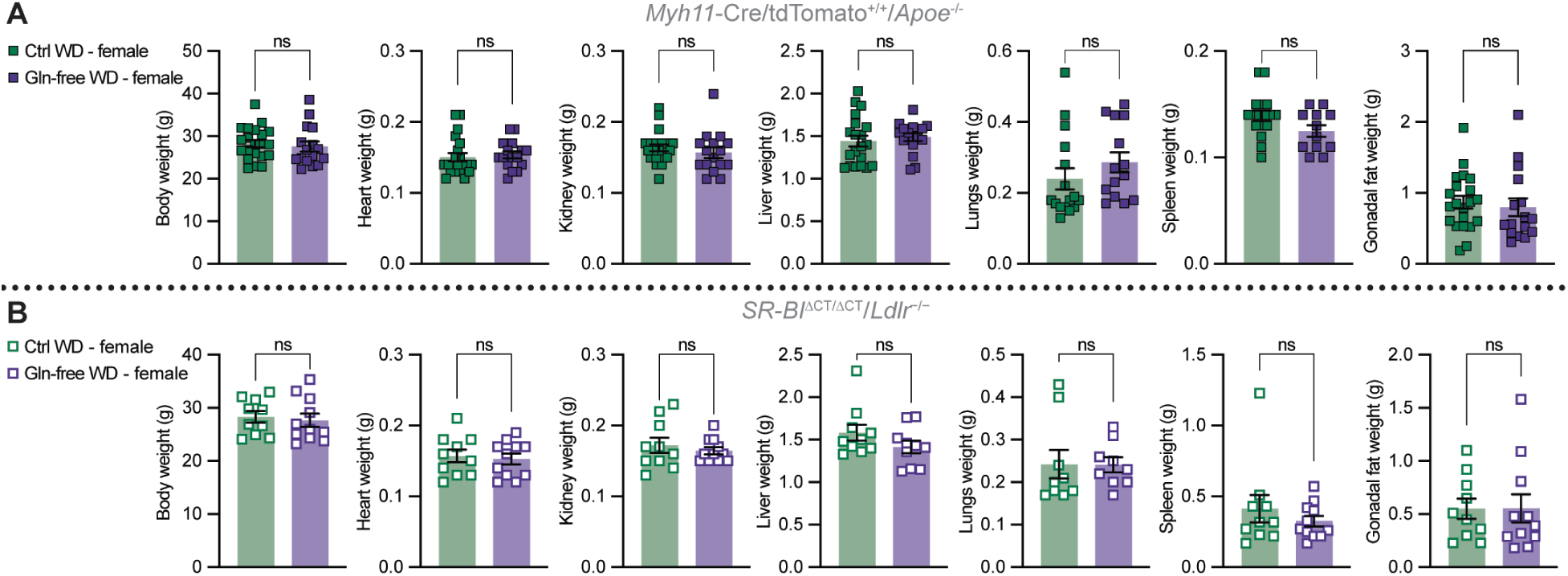
Body weight and organ weights in female mice fed a Western diet (WD) with or without glutamine for 18 and 30 weeks, related to Figure S9. (**A**) After 18 weeks of WD, no significant differences were observed between experimental groups in *Myh11*-Cre/tdTomato^+/+^/*Apoe*^-/-^ female mice (*N* ≥ 14 per group; mice from fig. S9, A to M). (**B**) Similarly, after 30 weeks of WD, no significant differences were found in *SR-BI*^ΔCT/ΔCT^/*Ldlr*^−/−^ female mice (*N* ≥ 10 per group; mice surviving the experiment from fig. S9, N to O). Graphs were analyzed using two-tailed unpaired *t*-test. Error bars represent the mean ± SEM. ns, not significant.

**Table S1.**
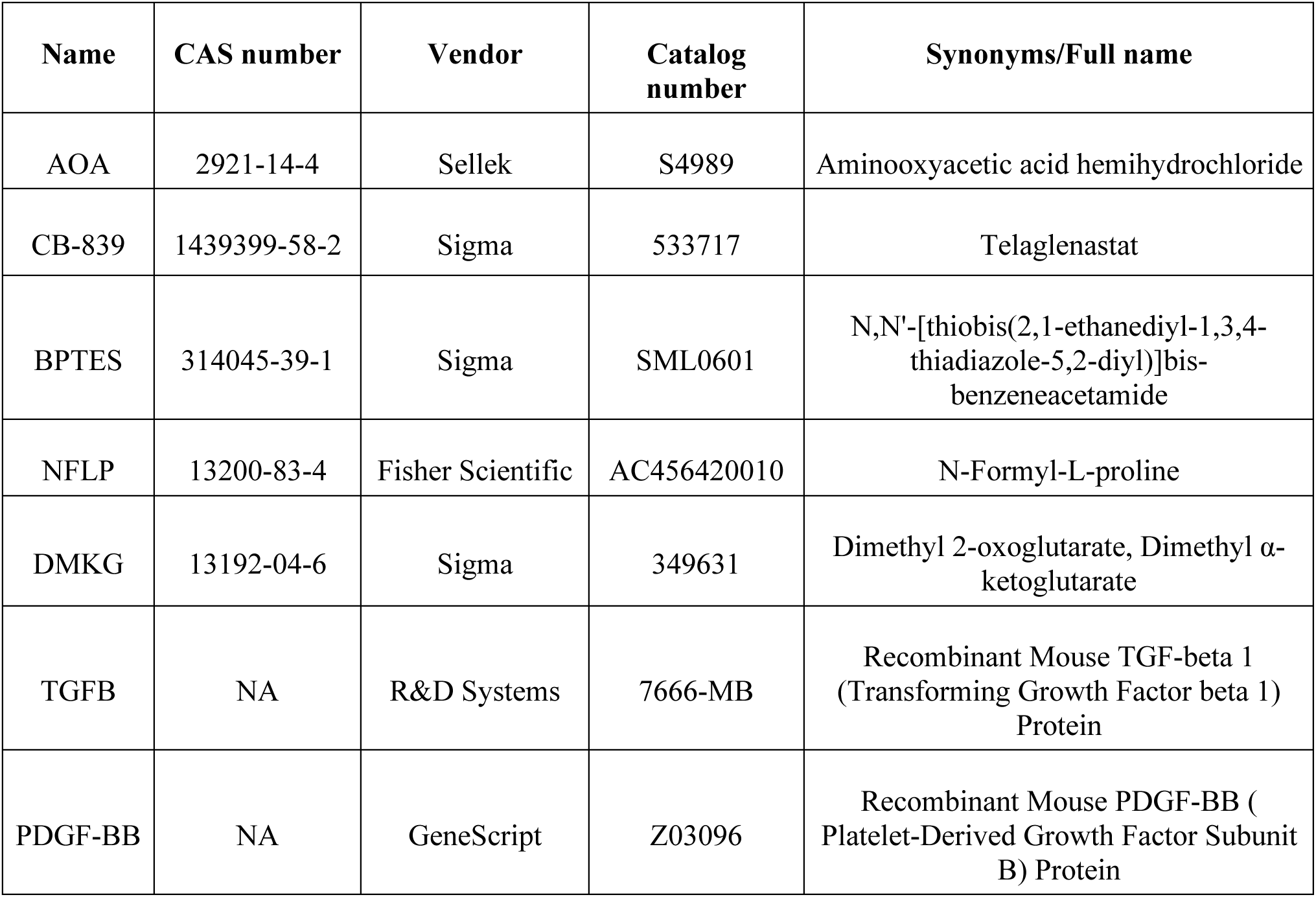
Treatment details for in vitro studies.

**Table S2.**
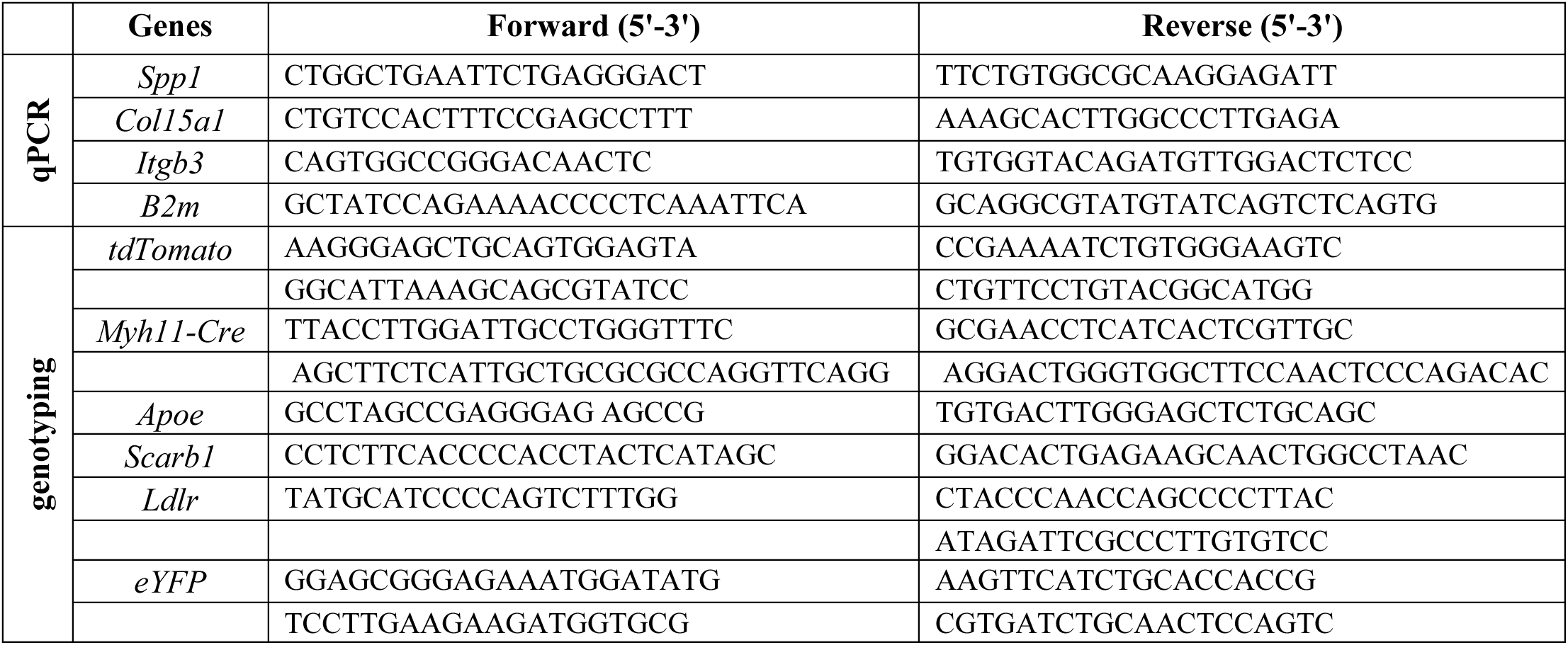
Primers for qPCR and genotyping.

**Data S1. Bulk RNA-seq analysis of M607 after PDGF/TGFB treatment for 24 hours, related to figure 1**.

Pathways enriched in Gene Set Enrichment Analysis (GSEA; clusterProfiler v4.10 ^82^) related to extracellular matrix organization (ECM), used for leading edge analysis for ECM-marker genes identification. FC: fold change. Gene ratio represents the proportion of genes annotated to a pathway (gene set) that reach a threshold for statistical significance (gene count) and in GSEA is equal to % tags from leading edge column.

**Data S2. Bulk RNA-seq and proteomics analysis of M607 at baseline (PDGF/TGFB vs vehicle) and after glutamine depletion (PDGF/TGFB-Gln vs PDGF/TGFB), related to Figure 1**.

Comprehensive analysis of common ECM gene and protein expression changes induced by PDGF+TGF treatment (baseline), and subset of baseline-changed ECM genes/proteins that is glutamine dependent (PDGF+TGF-Gln vs PDGF+TGF). Transcriptomic and proteomic datasets showing significant changes (adjusted *P*-value < 0.05).

**Data S3. Bulk RNA-seq super-pathway analysis of M437 treated with PDGF/TGFB and PDGF/TGFB-Gln at 0, 4, 24, 48, and 72 hours, related to Figure 2 and S4.**

Bulk RNA-seq data quantifying effect of presence or absence of glutamine in SMC-F transition over time. Only pathways meeting the filtering criteria are included. Values not meeting the thresholds are set to NA.

**Data S4. Bulk RNA-seq super-pathway analysis of M437 treated with PDGF/TGFB +/-Gln, DMKG, and AOA at 48 hours, related to Figure 2 and S4.**

Bulk RNA-seq data quantifying the effect of other treatments on SMC-F transitions after 48 hours. Only pathways meeting the filtering criteria are included. Values not meeting the thresholds are set to NA.

**Data S5. Super-pathway analysis of SMC-derived cells (tdTomato^+^) from scRNA-seq in male mice, related to Figure 3 and S6.**

Super-pathway analysis of differential expression data from comparing SMC transcriptional clusters. SMC-C1 was chosen as the reference point to normalize all other transcriptional clusters. Only pathways meeting the filtering criteria are included. Values not meeting the thresholds are set to NA.

**Data S6. Amino acid concentration in plasma from mice fed Ctrl WD or Gln-free WD for 18 weeks, related to Figure 4 and S8.**

Amino acid concentrations measured by mass spectrometry from plasma collected from mice at the end of the experiment via the cardiac puncture.

**Data S7. Amino acid concentration in plasma from mice fed Ctrl WD or Gln-free WD for 4 and 18 weeks, related to Figure 4 and S8.**

Amino acid concentrations measured by mass spectrometry from plasma collected mice via tail vein bleeding after 4 and 12 weeks of WD.

**Data S8. Bulk RNA-seq super-pathway analysis of hearts from male *SR-BI*^ΔCT/ΔCT^/*Ldlr*^-/-^mice fed either Gln-free WD or Ctrl WD for 30 weeks, related to Figure 5**.

Only pathways meeting the filtering criteria are included. Values not meeting the thresholds are set to NA.

